# Hepatic γδ NKT cells modulate liver-resident CD8+ T cells to attenuated malaria parasite vaccines

**DOI:** 10.64898/2026.08.21.746325

**Authors:** Rebecca C. Blyn, Aditi V. Kulkarni, Alexandra N. Donlan, Ashley A. Krakauer, Gina Jones, Sydney M. Nemphos, Noah Stegman, Emily G. Tanner, Nina Hertoghs, Katharine V. Schwedhelm, Stephen C. De Rosa, Kenneth D. Stuart, Cole Phalen, Lucas Graybuck, Peter J. Skene, Evan W. Newell, Suzanne M. McDermott, Nana K. Minkah

## Abstract

*Plasmodium* parasites develop in the liver and egress to infect red blood cells, causing malaria. Vaccines that generate hepatic CD8+ T cells eliminate liver-stage parasites and prevent disease, yet how these T cells are induced is incompletely understood. We report that in mice vaccinated with replication-competent genetically attenuated *Plasmodium* parasites, antagonism of γδ T cell function curtails protection. Vaccination expands hepatic IFNγ+ γδ NKT cells, and depletion of these cells abrogates hepatic CD8+ T cell responses. IFNγ+ γδ NKT cells are nearly undetectable in the blood at steady state but their frequencies in the periphery are significantly increased following vaccination, hinting at their utility as biomarkers of protection. To assess the relevance of these results in humans, we performed secondary analyses of peripheral blood samples from human clinical trial participants immunized with attenuated *Plasmodium* parasites (Trial registration: ClinicalTrials.gov NCT01994525). Flow cytometric and single cell transcriptomic characterization of γδ T cells in these samples unveil for the first time, increased frequency of activated Vδ2-γδ T cells and gene expression in cytotoxic, tissue-homing Vδ1+ γδ T cells as correlates of protection. Together, these data identify hepatic γδ T cells as targets for the improvement of tissue-resident CD8+ T cell responses against hepatotropic pathogens.

## INTRODUCTION

*Plasmodium spp.* parasites are the causative agents of malaria, a disease that kills 600,000 people annually. The *Plasmodium* lifecycle begins following the bite of an infected *Anopheles* mosquito, when it deposits sporozoites (**SPZs**) into the skin. These SPZs travel to the liver and infect hepatocytes as part of a clinically silent liver stage (**LS**). Following LS development (∼48 hours in mice and 5-7 days in humans), parasites egress into the bloodstream to infect red blood cells, resulting in clinical symptoms and malaria-associated mortality^1,2^. Blood-stage infection and clinical disease can be prevented by eliminating the pre-erythrocytic stages (SPZ and LS), making pre-erythrocytic stage vaccines attractive intervention strategies.

The most effective pre-erythrocytic stage vaccines are whole parasite vaccines (**WPVs**)^3^ in which the attenuation of live parasites causes them to arrest and die during the LS. These live-attenuated vaccines induce a multi-faceted immune response in the liver^4–7^ that can result in sterilizing immunity. The PfSPZ™ vaccine, composed of *P. falciparum* SPZs that have been irradiated to cause their early attenuation in the liver, has shown good efficacy in clinical trials^3^. However, recent studies show that late-arresting replication-competent genetically attenuated parasites (**LARC GAPs**), in which parasite genes critical for the completion of LS development have been removed, outperform the radiation-attenuated SPZ (**RAS**) vaccine, PfSPZ^TM^, and other early-arresting WPVs^8–11^. Nonetheless, LARC GAPs will need to be further optimized before they are deployed in mass immunization campaigns. Hepatic memory T cells, especially liver-resident memory CD8+ T cells (**T_RM_s**)^12,13^, are essential to WPV-engendered protection. A full understanding of the mechanisms that regulate T_RM_ differentiation, persistence, and recall will enable targeted improvement of these vaccines.

γδ T cells are unique T lymphocytes that express the γ and δ chains of the T cell receptor (**TCR**). They possess a variety of effector functions, can be rapidly activated, and are enriched in barrier and mucosal sites, positioning them as contributors to both innate and adaptive immunity^14–16^. γδ T cells have been identified as correlates of WPV-engendered protection in humans^7,10,11,17–21^; attenuated SPZ administration causes an increase in Vγ9Vδ2+ γδ T cell frequency and activation in the blood that often correlates with the development of immunity. In mice, recent work identified IL-4-producing splenic Vγ1+ cells as critical mediators of RAS-engendered protection^22^. These human and murine studies support the hypothesis that γδ T cells are not merely correlates of WPV-engendered protection but rather act as mechanistic orchestrators of the protective immune response to these early-arresting vaccines. Yet there are still several unknowns: i) do γδ T cells also modulate protective immunity in humans and mice vaccinated with highly protective LARC GAP vaccines, and what mechanisms do these γδ T cells employ? ii) In humans, γδ T cell populations differ greatly by location, with Vδ2+ γδ T cells enriched in the blood while Vδ2-(Vδ1/3+) γδ T cells are more abundant in tissues^23^. Live-attenuated parasite development is restricted to the liver, indicating that liver-associated immune cells may be especially critical in vaccine-induced protection. Are peripherally enriched γδ T cell subsets recruited into the liver to be activated or does immunization induce local hepatic γδ T cell subsets to regulate anti-*Plasmodium* immunity?

We leveraged well-established mouse models of *Plasmodium* LS immunization to show that γδ T cells are required for LARC GAP-induced sterile protection and that immunization specifically expands a population of liver-enriched Vγ1+Vδ6.3+ γδ natural killer (NK)T cells that produce IFNγ. Hepatic CD8+ T cells are essential to LARC GAP-induced protection^13^, and we now show that mice specifically depleted of Vγ1+ γδ T cells alone are unable to mount competent hepatic CD8+ T cell responses after LARC GAP immunization. Vγ1+Vδ6.3+ γδ NKT cells are enriched in the liver but can be found in the blood of LARC GAP-immunized mice, hinting at their potential to serve as peripheral biomarkers of WPV-engendered protection in humans. Human and mouse γδ T cells share conserved functions, yet differences in Vγ/Vδ usage between mice and human γδ T cells preclude the facile identification of a human correlate of the liver-enriched Vγ1+Vδ6.3+ γδ NKT cells identified in our study. We embarked on an unbiased bioinformatic analysis of single-cell transcriptomes generated from the blood of WPV-immunized humans to show that transcriptional responses in cytotoxic NK receptor-expressing Vδ1+ γδ T cells are associated with the development of protective immunity against *Plasmodium* infection. Additionally, flow cytometric analyses of these samples revealed for the first time increased frequencies of activated, NK receptor-expressing Vδ2-cells as a correlate of vaccine-engendered protection. Together, these findings implicate liver-enriched γδ T cell subsets as regulators of the protective hepatic CD8+ T cell response to live-attenuated malaria parasite vaccines.

## RESULTS

### γδ T cells are essential for LARC GAP-induced protective immunity

To characterize the role of γδ T cells in LARC GAP-engendered protection, we utilized a partially protective immunization regimen^13^ in which wildtype C57BL/6J (**B6**) mice, TCRδ^−/-^mice, or B6 mice treated with a γδ T cell-inhibiting antibody^24^ (**α-TCRγδ**) were first immunized twice with LARC GAP (*Py fabb/f^−^*^25^) SPZs or debris isolated from the salivary glands of uninfected mosquitos (**mock**) and then challenged approximately 40 days later with luciferase-expressing wildtype (**WT**) parasites (***Py luc***^26^) (**Fig. 1A**). 42 hours post-challenge, we utilized *in vivo* bioluminescent imaging to assess parasite burden in the livers of B6 and TCRδ^−/-^mice. In comparison to mock-immunized mice, LARC GAP immunization resulted in a ∼170-fold reduction in LS burden in B6 mice (**Fig. 1B**). In contrast, we observed a ∼9-fold decrease in LS burden between mock-immunized and LARC GAP-immunized TCRδ^−/-^mice, indicating that parasite control in immunized mice is defective in the absence of γδ T cells (**Fig. 1B**). LS burden did not differ between mock-immunized B6 or TCRδ^−/-^mice.

**Figure 1:**
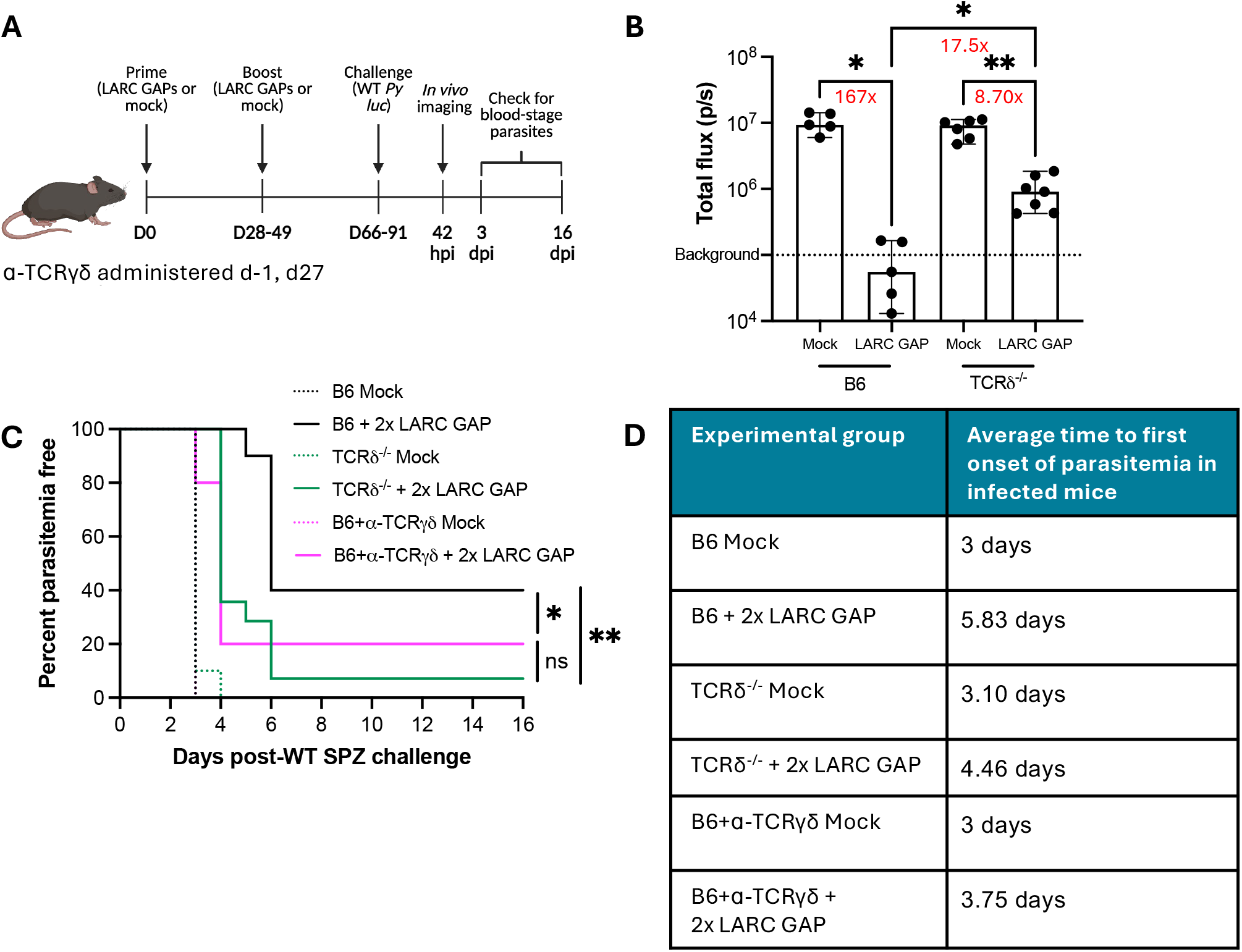
γδ T cells regulate the adaptive immune response to LARC GAP immunization. **A)** Mice were immunized twice with 50k LARC GAP SPZ or mock. Following the second immunization, all mice were challenged with 10k *Py luc* SPZ. One cohort of B6 and TCRδ^−/-^mice underwent *in vivo* bioluminescent imaging 42 hours post-infection (hpi) to assess parasite burden in the liver. Daily blood smears were collected to assess time to blood-stage infection. **B)** Total flux in mock-vs. LARC GAP-immunized B6 and TCRδ^−/-^ mice at 42 hpi. Numbers in red show fold-change differences in luminescence between selected groups. **C)** Time to blood-stage infection/patent parasitemia in mock-vs. LARC GAP-immunized B6 mice, TCRδ^−/-^ mice, and B6 mice treated with ɑ-TCRγδ at d-1 and 27. **D)** Table showing average time to blood-stage infection for infected mice in each group. Number of mice per group in **(C-D)**: B6 Mock – 8, B6 + 2x LARC GAP – 10, TCRδ^−/-^ Mock – 10, TCRδ^−/-^ + 2x LARC GAP – 14, B6+ɑ-TCRγδ Mock – 5, B6+ɑ-TCRγδ + 2x LARC GAP – 5. All bar plots are shown as the median values ± 95% confidence interval. Each dot represents one mouse. For *in vivo* imaging results, data were tested for significant outliers via Grubb’s test and compared via the Brown Forsythe and Welch ANOVA test followed by Dunnett’s T3 multiple comparisons test. Survival curves were compared using the Mantel-Cox test. All data come from at least two independent experimental replicates except for **(B)** and B6+ɑ-TCRγδ in **(C)**, which were performed once. *p<0.05, **p<0.01

Next, we assessed the efficacy of the LARC GAP vaccine in preventing the onset of blood-stage infection in B6 and γδ T cell-deficient mice. Starting at three days post-challenge, we assessed parasite presence in the blood (parasitemia). Two doses of LARC GAPs provided sterile immunity in 40% of B6 mice, a significant increase in protection compared to mock-immunized B6 mice, which all developed blood-stage infections (**Fig. 1C**). Furthermore, for the 60% of LARC GAP-immunized B6 mice that did not exhibit sterilizing immunity, we observed a nearly three-day delay in the time to onset of blood-stage infection as compared to mock-immunized mice (**Fig. 1D**). Notably, only one LARC GAP-immunized TCRδ^−/-^ mouse out of 14 was sterilely protected (**Fig. 1C**). Additionally, LARC GAP-immunized TCRδ^−/-^ mice that developed blood-stage infections did so sooner than immunized B6 counterparts (**Fig. 1D**). This impairment in LARC GAP-engendered protection in TCRδ^−/-^ mice was also observed in B6 mice treated with α-TCRγδ during immunization (**Fig. 1C-D**), reinforcing the importance of γδ T cells as modulators of LARC GAP-engendered protection.

### Diverse populations of γδ T cells respond early to LARC GAP immunization

To characterize the γδ T cells induced by LARC GAP immunization, we immunized B6 mice with LARC GAP SPZs or uninfected salivary glands and isolated and characterized leukocytes from the livers, spleens, and liver-draining lymph nodes (**LDLNs**) of each animal via cytometry by time of flight (**CyTOF)** over four timepoints (4 hours and days 3, 7, and 14 post-immunization). Phenograph clusters were generated via unsupervised clustering based on protein expression to show that LARC GAP immunization results in 28 clusters of γδ T cells (**Fig. 2A**). Populations shifted rapidly over time and across anatomical sites (**Fig. 2B**). At 4 hours post-immunization (**hpi**), γδ T cell clusters in mock-and LARC GAP-immunized mice in all organs were largely the same, although the clusters in the liver differed from those in the spleen and LDLNs, highlighting anatomical differences at baseline. By day 3 post-immunization, LARC GAP-induced changes to γδ T cell populations became evident in all organs. Interestingly, while clusters in the liver and spleen in LARC GAP-immunized mice showed remarkable similarity despite differences at 4 hpi, γδ T cell clusters in the LDLNs of LARC GAP-immunized mice remained unique. Rapid changes again occurred in all three organs between days 3 and 7 post-vaccination, with hepatic γδ T cell clusters becoming more distinct from the populations in the spleen and LDLNs. Fewer changes occurred in the spleen and LDLNs between days 7 and 14 post-immunization, but the liver still showed evidence of key cluster alterations occurring and had distinct population structures compared to the secondary lymphoid organs (**Fig. 2B**).

**Figure 2:**
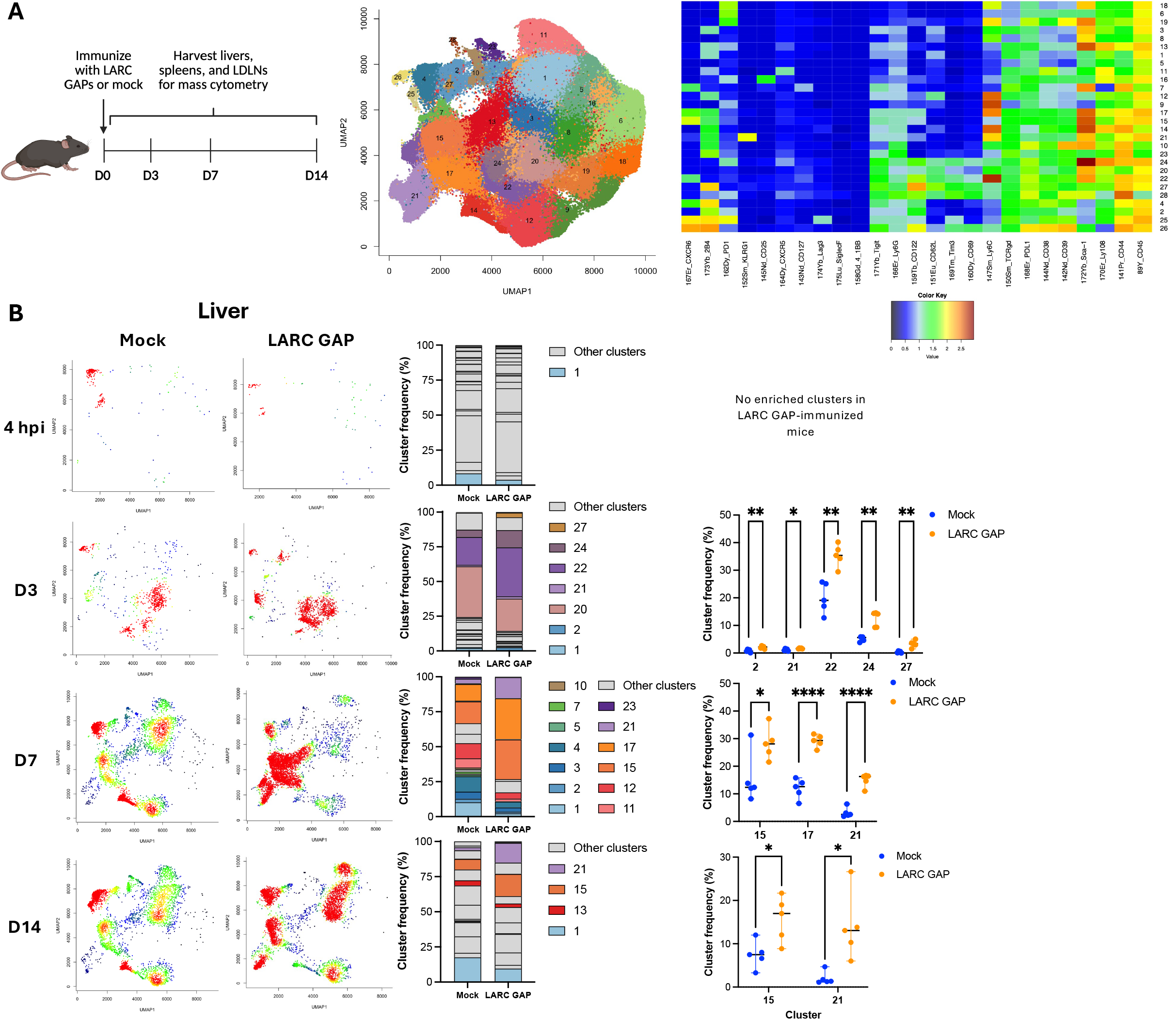
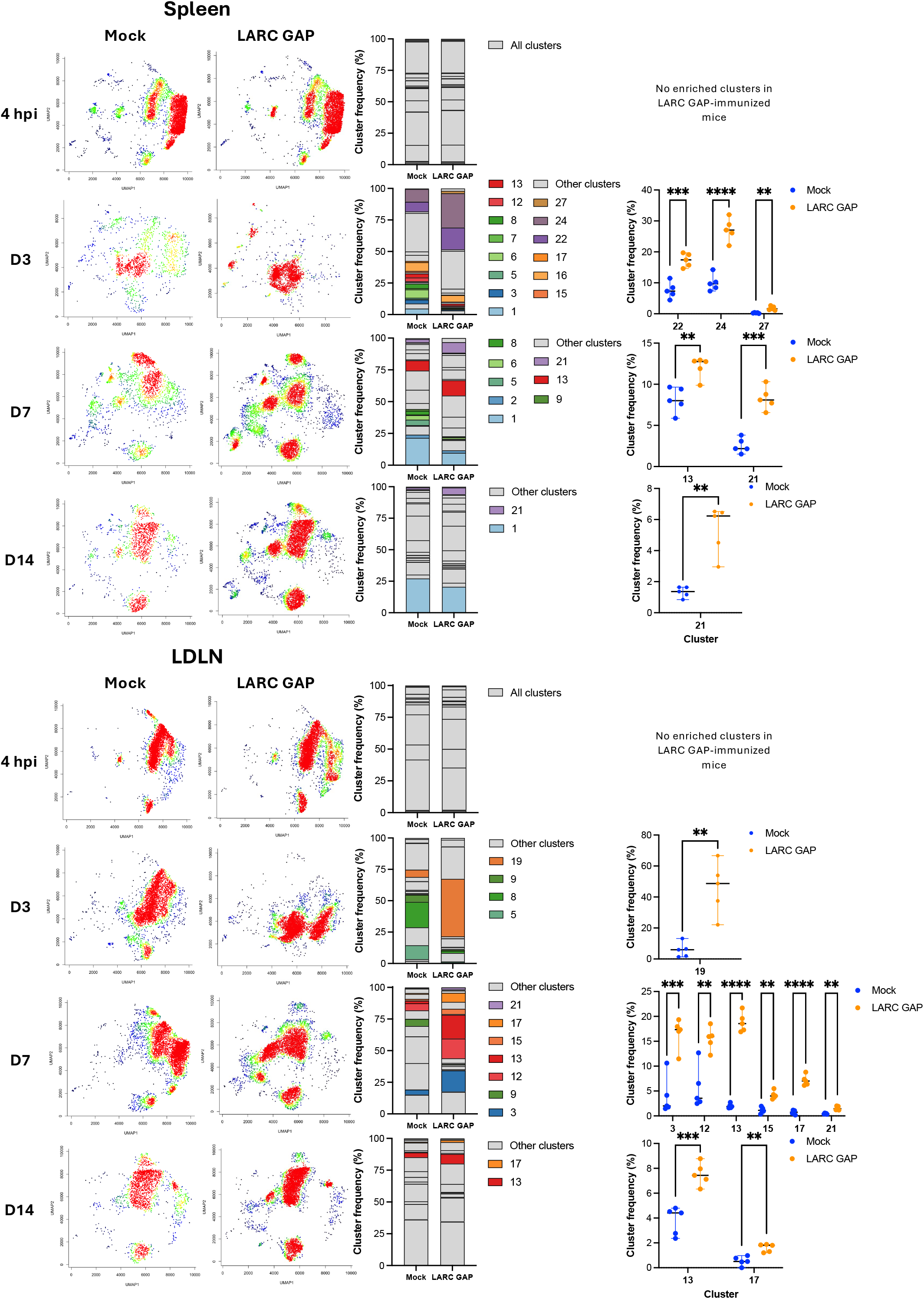
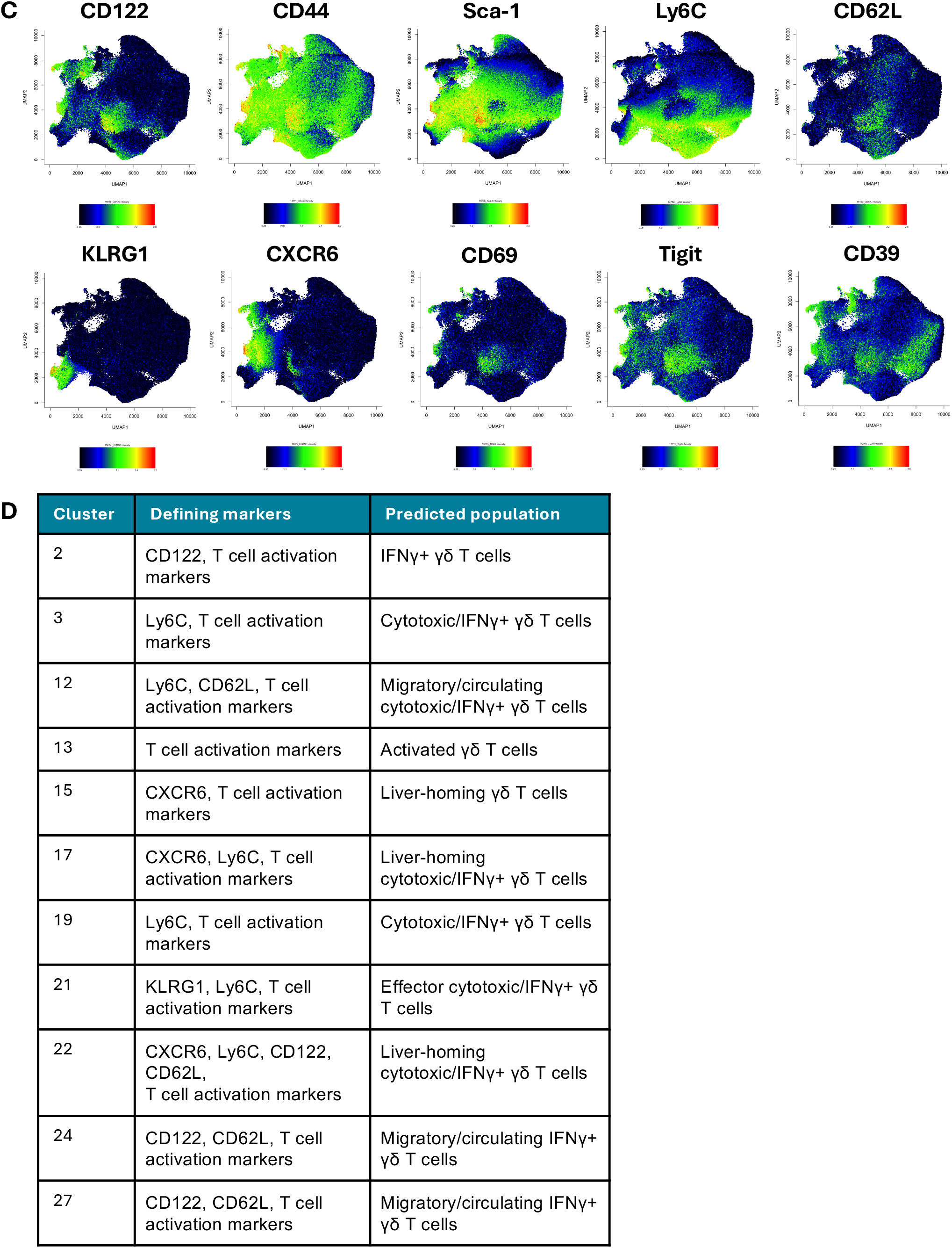
LARC GAP vaccination induces dynamic and heterogenous populations of γδ T cells across anatomical sites early post-vaccination. **A)** B6 mice were immunized with 100k LARC GAP SPZ or mock. Organs were harvested from all mice at 4 hours and days 3, 7, and 14 post-immunization for mass cytometry analysis. A phenograph showing the 28 γδ T cell clusters was generated utilizing protein expression, as shown in the heatmap. A list of all clusters and their protein markers can be found in **Table S1**. **B)** (Left) Cluster maps generated for each condition, timepoint, and organ. Red indicates the highest density of cells while blue indicates the lowest density. Each dot is a single cell. (Middle) Stacked bar plots showing expression of all clusters at each timepoint. Only clusters that had significantly increased frequency in mock-or LARC GAP-immunized mice are shown in color. (Right) Plots showing frequency of clusters enriched in LARC GAP-immunized mice compared to mock-immunized mice at each timepoint. A list of clusters enriched in mock-immunized mice compared to LARC GAP-immunized mice can be found in **Table S2. C)** Phenographs depicting protein expression. Red indicates the highest density of cells while blue indicates the lowest density. **D)** Table showing protein expression and predicted populations for clusters enriched in LARC GAP-immunized mice. For stacked bar plots showing cluster frequency, values were averaged across all mice. For plots showing frequency of selected clusters, frequencies are shown as the median ± 95% confidence interval, and each dot is one mouse. Frequencies were compared using Welch’s t-test. Data represent one replicate with 5 mice per group. *p<0.05, **p<0.01, ***p<0.001, ****p<0.0001

We next utilized published datasets^27–32^ to classify the γδ T cell clusters identified in our CyTOF data, which revealed an expansion of IFNγ+ and cytotoxic effector γδ T cells in response to the LARC GAP vaccine. In the liver, clusters that were more prevalent in LARC GAP-immunized mice compared to mock-immunized mice appeared at day 3 post-immunization. These day 3 clusters included 2, 22, 24, and 27, which expressed canonical T cell activation markers (e.g. CD69, Tigit, CD38, CD39) alongside CD122, a known marker of IFNγ-expressing γδ T cells^27^ (**Fig. 2B-D**). Cluster 21 was also enriched in the livers of LARC GAP-immunized mice and showed increased expression of KLRG1, marking it as a possible effector γδ T cell population^28^ (**Fig. 2B-D**). Additionally, clusters 21 and 22 expressed Ly6C, which has been identified as a potential marker of cytotoxic and IFNγ-producing γδ T cells^29,30^ (**Fig. 2C, D**). By day 7 post-immunization, cluster proportions in the liver shifted dramatically, with livers from LARC GAP-immunized mice demonstrating significantly higher frequencies of clusters 15, 17, and 21 compared to livers from mock-immunized mice (**Fig. 2B**). Furthermore, these three clusters together accounted for 75% of all hepatic γδ T cells in LARC GAP-immunized mice (**Fig. 2B**). Interestingly, clusters 15 and 17 expressed CXCR6, a liver-homing/residency marker^33^ (**Fig. 2C**). Cluster 15 also exhibited high expression of CD44 and Sca1 without CD122, which may mark it as an IL-17-expressing γδ T cell population^31^ (**Fig. 2C, D**). However, γδ T cells in the liver are known to express high levels of CD44 even at baseline^32^ and Sca1 may also serve as an activation marker, making it difficult to truly identify IL-17+ γδ T cells. Similarly to cluster 21, cluster 17 showed high expression of Ly6C and thus may represent cytotoxic IFNγ+ γδ T cells (**Fig. 2C, D**). γδ T cell subsets in the liver continued to shift between day 7 and 14 post-immunization, with clusters 15 and 21 maintaining elevated frequencies in LARC GAP-immunized mice (**Fig. 2B**). Overall, the population shifts between days 3, 7, and 14 post-immunization indicate that IFNγ+ and cytotoxic effector γδ T cells dominate the hepatic response to LARC GAP vaccination.

We also identified alterations to γδ T cell populations in the spleen and LDLNs in response to LARC GAP vaccination. While day 3 splenic clusters enriched in LARC GAP-immunized mice were similar to those in the liver (**Fig. 2B**), LDLNs from LARC GAP-immunized mice showed increased frequency of cluster 19, which had high Ly6C expression alongside other activation markers (**Fig. 2B, C**). Clusters shifted in the secondary lymphoid organs by day 7, with more shared enriched clusters across all organs in LARC GAP-immunized mice (**Fig. 2B**). Uniquely in the LDLNs and spleen, LARC GAP vaccination resulted in an increase in the frequency of cluster 13, which expressed high levels of Sca1, Ly108, and CD44 (**Fig. 2B, C**). In the LDLNs alone, clusters 3 and 12 were significantly enriched in LARC GAP-immunized mice, with both clusters expressing Ly6C and cluster 12 also expressing CD62L, possibly marking it as a migratory/circulating γδ T cell population^28^ (**Fig. 2B-D**). γδ T cell populations in the spleen and LDLNs from LARC GAP-immunized mice appeared largely like those in mock-immunized mice and to each other by day 14 post-immunization (**Fig. 2B**). However, LDLNs from LARC GAP-immunized mice showed continued elevated abundance of clusters 13 and 17, and spleens from LARC GAP-immunized mice displayed elevated frequency of cluster 21, similar to the liver (**Fig. 2B**). Thus, our high-parameter CyTOF analyses indicate that γδ T cell population-level changes occur temporally and spatially early post-immunization, with a tendency in all organs towards increased proportions of predicted IFNγ+ effector subsets in LARC GAP-immunized mice.

### Hepatic γδ NKT cells expand following vaccination with LARC GAPs

To further define the specific γδ T cell subsets induced by LARC GAPs, we immunized B6 mice with LARC GAP SPZs or uninfected salivary glands and isolated leukocytes from the livers, spleens, and LDLNs at day 4 post-immunization for flow cytometric analysis (**Fig. 3A**). We observed a significant 2-fold increase in the number of γδ T cells in the liver after vaccination but saw no significant change in the number of γδ T cells in the spleen or LDLNs (**Fig. 3B**). This was despite finding significant alterations to γδ T cells in the secondary lymphoid organs via our CyTOF analyses, suggesting that these changes may have been specifically to γδ T cell cellular states without altering the number of cells present. We additionally found that the numbers of hepatic γδ T cells that expressed CD62L, CD69, and CD122 were increased following LARC GAP vaccination, consistent with our mass cytometry findings (**Fig. S1A-C**). Vaccination also led to a substantial increase in hepatic γδ T cells that expressed the activation markers CD160 and ICOS and an increase in proliferating Ki67+ γδ T cells (**Fig. S1D-F**).

**Figure 3:**
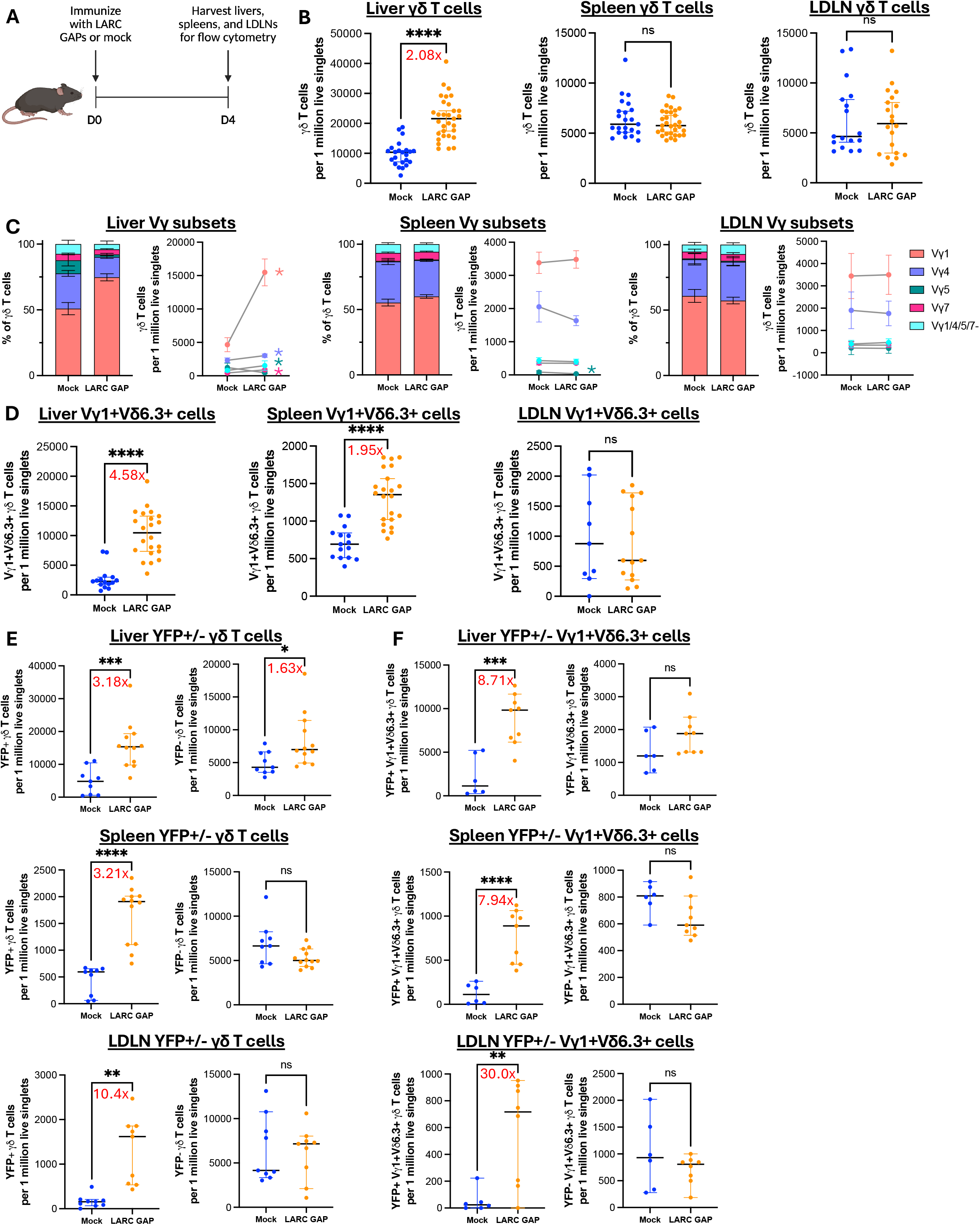
IFNγ+ Vγ1+Vδ6.3+ T cells are induced early in the liver following LARC GAP immunization. **A)** B6 and GREAT mice were immunized with LARC GAP SPZ or mock. Organs were harvested from all mice at day 4 post-immunization. **B)** Number of γδ T cells per 1 million live singlet lymphocytes. **C)** Frequency of Vγ chain expression in γδ T cells and Vγ chain populations expressed per 1 million live singlet lymphocytes. Asterisks are shown next to populations that differ significantly between mock and LARC GAP-immunized mice. Liver: Vγ1: p<0.0001; Vγ4: p<0.01; Vγ5: p<0.05; Vγ7: p<0.0001. Spleen: Vγ5: p<0.05. **D)** Number of Vγ1+Vδ6.3+ cells per 1 million live singlet lymphocytes. **E)** Number of YFP+ and YFP-γδ T cells per 1 million live singlet lymphocytes. **F)** Number of YFP+ and YFP-Vγ1+Vδ6.3+ cells per 1 million live singlet lymphocytes. All dot plots show the median ± 95% confidence interval, and each dot represents one mouse. Red numbers above each plot show the fold-change difference between the selected groups. For stacked bar plots showing Vγ chain frequency, plots show normalized median values ± 95% confidence intervals. For paired dot plots showing number of Vγ-expressing cells, plots show median values ± 95% confidence intervals. Data were compared using Welch’s t-test. All data come from at least two independent experimental replicates. *p<0.05, **p<0.01, ***p<0.001, ****p<0.0001

To further classify γδ T cell subsets, we utilized antibodies that target the Vγ chain of the γδ TCR. Although the frequencies of different Vγ-expressing γδ T cells were largely unaltered by vaccination in the spleen and LDLNs, we observed a strong increase in the frequency and number of Vγ1+ γδ T cells in the liver at day 4 post-immunization, with ∼75% of all γδ T cells expressing the Vγ1 chain in LARC GAP-immunized mice (**Fig. 3C**, Heilig and Tonegawa nomenclature^34^). The vast majority of these Vγ1+ cells co-expressed the Vδ6.3 chain, marking them as γδ NKT cells^32,35^ (**Fig. 3D, Fig. S2**). While the strongest induction of these γδ NKT cells by LARC GAP vaccination was seen in the liver (with over 4.5-fold expansion compared to mock-immunized mice), γδ NKT cells were also enriched in the spleens of LARC GAP-immunized mice by almost 2-fold (**Fig. 3D**). Thus, our data indicate that γδ NKT cells are uniquely induced by LARC GAP vaccination, with the major impact occurring in the liver and to a lesser extent in the spleen.

### LARC GAP vaccination induces IFNγ expression in hepatic γδ NKT cells

Our CyTOF data suggested that LARC GAP-induced γδ T cell populations were likely to be IFNγ-producing. Hepatic γδ NKT cells express IFNγ upon activation^32,36^, and so we immunized interferon-<u>g</u>amma reporter with <u>e</u>ndogenous poly<u>A</u> transcript (**GREAT)** mice that express YFP under the control of the IFNγ promoter^37^ and examined YFP expression by γδ T cells four days later. We observed a significant ∼3.2-fold increase in bulk YFP+ γδ T cells in the liver (**Fig. 3E**). While YFP-cells were also significantly enriched in LARC GAP-immunized livers by ∼1.6-fold, they were a minority compared to the YFP+ population (**Fig. 3E**). YFP+ γδ T cells also expanded in the spleens and LDLNs of LARC GAP-immunized mice by ∼3.2-fold and ∼10.4-fold respectively, although YFP-γδ T cells in these organs were the dominant population (**Fig. 3E**). Intracellular staining of IFNγ showed a similar increase in IFNγ expression in hepatic γδ T cells following vaccination (**Fig. S1G**). Vγ4+ and Vγ6+ cells showed low IFNγ expression as expected^38^ (**Fig. S3**), demonstrating that background levels of YFP expression in GREAT mice were low.

To confirm that γδ NKT cells expressed IFNγ, we examined YFP expression in the Vγ1+Vδ6.3+ population. While YFP-Vγ1+Vδ6.3+ cells showed no significant expansion in the liver following vaccination, YFP+ Vγ1+Vδ6.3+ cells increased almost 9-fold (**Fig. 3F**). Additionally, YFP+ Vγ1+Vδ6.3+ cells outnumbered YFP+ Vγ1+Vδ6.3-cells in the livers of LARC GAP-immunized mice by ∼5-fold, and over 80% of Vγ1+Vδ6.3+ cells expressed YFP as compared to ∼40% of Vγ1+Vδ6.3-cells (**Fig. 3F, Fig. S2**). Furthermore, this enrichment of YFP+ Vγ1+Vδ6.3+ cells was also observed in the spleen and LDLNs, although the number of YFP+ Vγ1+Vδ6.3+ cells in the liver was significantly higher (**Fig. 3F**). Altogether, these results expand on our CyTOF experiments to show that LARC GAP vaccination induces an early expansion of IFNγ-expressing γδ NKT cells across anatomical sites, with the dominant response occurring in the liver.

### Vγ1+ γδ T cells regulate LARC GAP-induced hepatic CD8+ T cell responses

Given that LARC GAP-engendered protective immunity is dependent on the generation of anti-*Plasmodium* hepatic CD8+ T cells^13^, we hypothesized that γδ T cells may regulate the hepatic CD8+ T cell response to vaccination. To address this, we treated B6 mice with an IgG antibody control or α-TCRγδ monoclonal antibody and immunized all animals with LARC GAP SPZ or uninfected salivary glands. Seven days later, we assessed CD8+ and CD4+ T cell frequencies in the liver and spleen (**Fig. 4A, Fig. S4A-B**). In B6 mice treated with the IgG control antibody, LARC GAP immunization resulted in a ∼2.6-fold increase in bulk hepatic CD8+ T cells (**Fig. 4B**) and ∼4.6-fold increase in activated hepatic CD8+ T cells (**Fig. 4C**) as compared to mock-immunized mice. However, in B6 mice that were treated with α-TCRγδ, no significant increase in hepatic bulk or activated CD8+ T cells was observed following LARC GAP immunization (**Fig. 4B, C**). No impact of γδ T cell inhibition was seen on either CD8+ T cells in the spleens or on CD4+ T cells in either organ (**Fig. 4B-C, Fig. S4A-B**), indicating that the regulatory role of γδ T cells is specific to the induction of hepatic CD8+ T cells.

**Figure 4:**
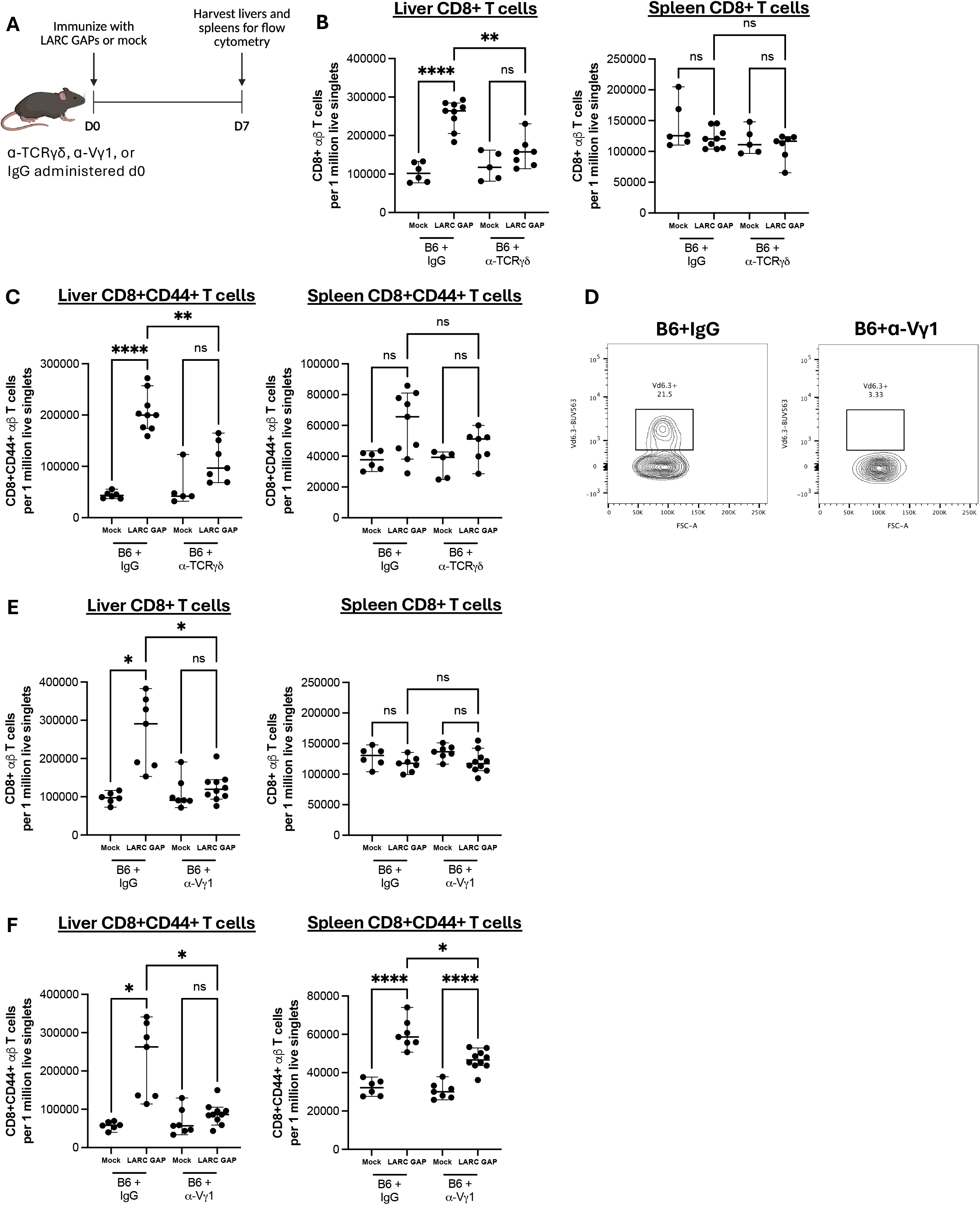
Vγ1+ T cells regulate the adaptive hepatic CD8+ T cell response to LARC GAPs. **A)** Mice were treated with an IgG control antibody, ɑ-TCRγδ (**B,C**), or ɑ-Vγ1 (**E,F**) antibody on day 0 prior to mock or LARC GAP immunization. 7 days later, organs were harvested for flow cytometry. **B, E)** Number of CD8+ T cells per 1 million live singlet lymphocytes in the liver and spleen. **C, F)** Number of CD8+CD44+ T cells per 1 million live singlet lymphocytes in the liver and spleen. **D)** Sample flow cytometry plot showing population of Vδ6.3+ cells in the livers of naïve TCRδ-eGFP mice treated with IgG or ɑ-Vγ1 24 hours before harvest. Cells were gated on CD3+TCRβ-live singlet lymphocytes. All plots show the median ± 95% confidence interval. Each dot represents one mouse. The Brown-Forsythe and Welch ANOVA test followed by Dunnett’s T3 multiple comparisons test were used to statistically analyze the data. All data come from at least two independent experimental replicates except for (**D)**, which was performed once. *p<0.05, **p<0.01, ****p<0.0001

We next treated B6 mice with IgG or α-Vγ1 (a monoclonal antibody to deplete Vγ1+ cells) followed by immunization with LARC GAP SPZ or uninfected salivary glands and examined T cell frequencies in the liver and spleen seven days later (**Fig. 4A, Fig. S4C-D**). Depletion of Vγ1+ cells eliminated the hepatic Vγ1+Vδ6.3+ cell population (**Fig. 4D**) and reduced total hepatic γδ T cells (**Fig. S5**). B6 mice treated with IgG showed a ∼3-fold increase in bulk hepatic CD8+ T cells and ∼4.5-fold increase in activated hepatic CD8+ T cells following LARC GAP immunization (**Fig. 4E, F**). Strikingly, depletion of Vγ1+ cells alone abrogated the hepatic CD8+ T cell response, replicating the defect observed when utilizing the pan-γδ T cell inhibiting antibody (**Fig. 4E, F**). In this set of experiments, we also found that activated CD8+ T cells increased slightly in the spleens of LARC GAP-immunized mice treated with IgG or α-Vγ1 by ∼1.8-fold and ∼1.6-fold, respectively (**Fig. 4F**). Additionally, Vγ1 depletion caused a slight ∼20% decrease in the number of activated CD8+ T cells in the spleens of LARC GAP-immunized mice compared to IgG-treated mice but did not fully eliminate the vaccine-induced response (**Fig. 4F**). CD4+ T cells in the liver and spleen were unaffected by the loss of Vγ1+ cells (**Fig. S4C, D**). Overall, these results reinforce that amongst hepatic γδ T cells, Vγ1+ cells are the subset responsible for modulating the anti-*Plasmodium* effector CD8+ T cell response in the liver.

### IFNγ-expressing γδ NKT cells can be tracked in the blood post-immunization

Our results show that IFNγ+ γδ NKT cells expand in the liver post-vaccination and regulate LARC GAP-induced adaptive immunity, indicating that these cells may be useful vaccine-induced biomarkers. However, the human liver is inaccessible for immune analysis during clinical trials, making it difficult to characterize hepatic γδ T cells in vaccinated individuals. Tissue-resident and tissue-enriched T cells can traffic out of tissues into the periphery^39^. We therefore set out to determine if IFNγ+ γδ NKT cells could be identified in the blood of immunized mice (**Fig. 5A**). On day 4 post-immunization, we observed a significant ∼1.7-fold increase in the number of all γδ T cells in the blood of LARC GAP-immunized mice (**Fig. 5B**) and a ∼2.9-fold increase in the number of Vγ1+ cells (**Fig. 5C**). Additionally, the number of Vγ1+Vδ6.3+ cells in the blood of LARC GAP-immunized mice was increased nearly 4-fold (**Fig. 5D**). These increases were even more striking when we enumerated the number of IFNγ-producing γδ T cells; YFP+ γδ T cells showed a robust ∼11.5-fold increase in the blood post-LARC GAP immunization (**Fig. 5E**), while YFP+ Vγ1+Vδ6.3+ cells increased by over 18-fold in LARC GAP-immunized mice (**Fig. 5F**). No changes in YFP-γδ T cell populations were observed in the blood (**Fig. 5E-F**).

**Figure 5:**
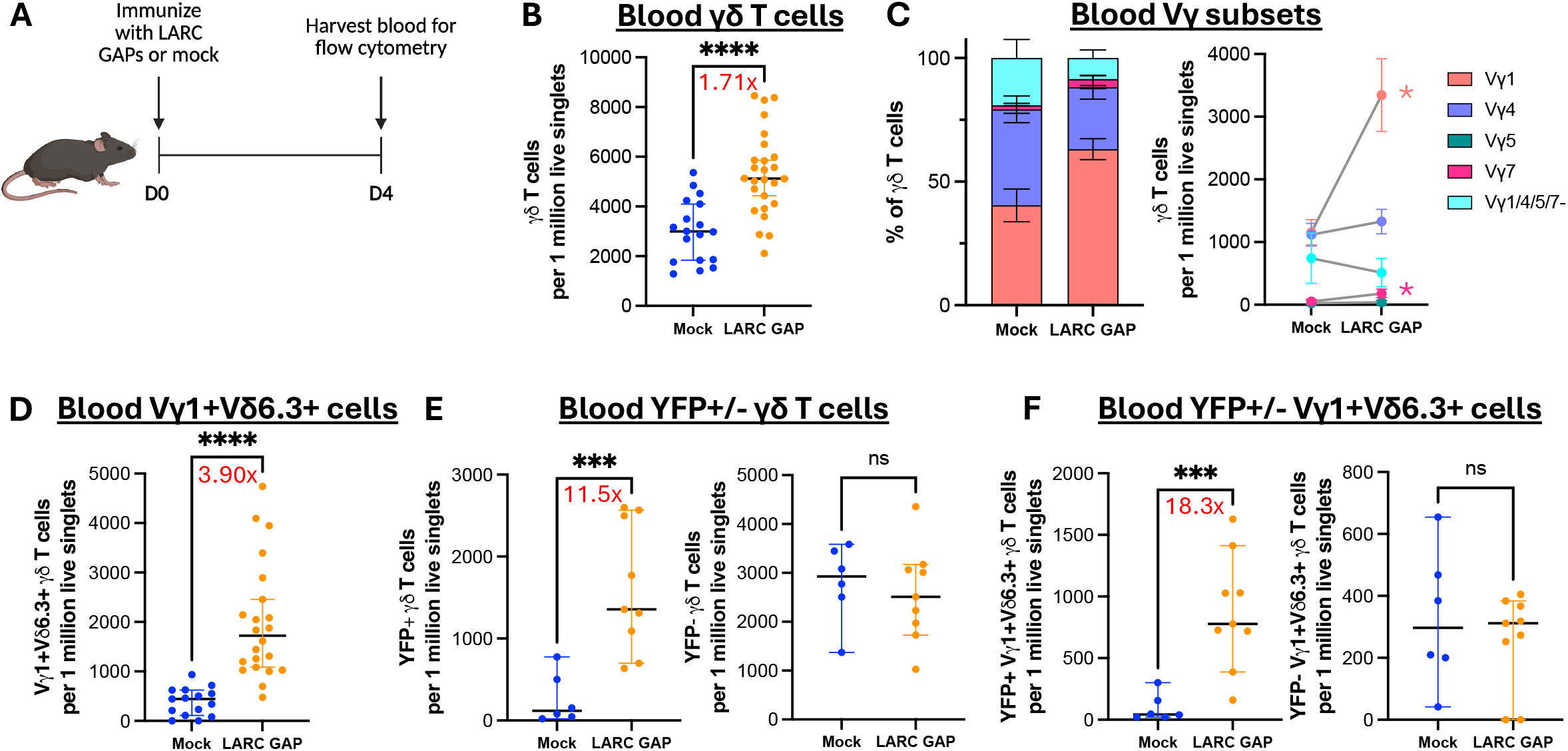
IFNγ+ Vγ1+Vδ6.3+ T cells are induced early in the blood following LARC GAP immunization. **A)** B6 and GREAT mice were immunized with LARC GAP SPZ or mock. Blood was harvested at day 4 post-immunization. **B)** Number of γδ T cells per 1 million live singlet lymphocytes. **C)** Frequency of Vγ chain expression in γδ T cells and Vγ chain populations expressed per 1 million live singlet lymphocytes. Asterisks are shown next to populations that differ significantly between mock and LARC GAP-immunized mice. Vγ1: p<0.0001; Vγ7: p<0.01. **D)** Number of Vγ1+Vδ6.3+ cells per 1 million live singlet lymphocytes. **E)** Number of YFP+ and YFP-γδ T cells per 1 million live singlet lymphocytes. **F)** Number of YFP+ and YFP-Vγ1+Vδ6.3+ cells per 1 million live singlet lymphocytes. All dot plots show the median ± 95% confidence interval. Each dot represents one mouse. Red numbers above each plot show the fold-change difference between the selected groups. For stacked bar plots showing Vγ chain frequency, plots show normalized median values ± 95% confidence intervals. For paired dot plots showing number of Vγ-expressing cells, plots show median values ± 95% confidence intervals. Data were compared using Welch’s t-test. All data come from at least two independent experimental replicates. ***p<0.001, ****p<0.0001

### Changes to human T and NKT-like cells upon attenuated SPZ immunization and Controlled Human Malaria Infection

Our data thus far indicate that liver-enriched Vγ1+Vδ6.3+ γδ NKT cells modulate the development of hepatic CD8+ T cell responses to *Plasmodium* infection of the murine liver. Frequencies of these γδ NKT cells are increased in the blood of LARC GAP-immunized animals, albeit as a minority population of all γδ T cells in the blood. We hypothesized that if WPVs also activate liver-enriched γδ T cells, then human equivalents of Vγ1+Vδ6.3+ γδ T cells in the blood might serve as useful correlates of WPV-engendered protection. Human malaria clinical trial studies have often identified Vγ9+Vδ2+ γδ T cells as a correlate of protection, yet given their enrichment in the blood and not tissues^23^, we reasoned that Vγ9+Vδ2+ γδ T cells might not be the human correlates of the murine cells identified. Murine and human γδ T cells are distinctly classified by their Vγ or Vδ chain respectively, making direct comparison of these subsets difficult. To identify potential human equivalents of WPV-induced Vγ1+Vδ6.3+ γδ T cells, we analyzed a single-cell transcriptome dataset generated from peripheral blood mononuclear cells (**PBMCs**) isolated from PfSPZ^TM^-vaccinated individuals in cohort 1 of the Immunization by Mosquito bite with Radiation Attenuated SPZ (**IMRAS**) trial^40^ (**Fig. 6A**). In this trial, malaria-naïve volunteers received five doses of PfSPZ^TM^ via mosquito bite and protection was assessed following Controlled Human Malaria Infection (**CHMI**) (**Fig. 6A**). With 55% of the vaccinees exhibiting sterile protection against SPZ challenge, cohort 1 offers a unique opportunity to determine if a cytotoxic, tissue-enriched human γδ T cell correlates with the development of protection. Leukapheresis samples were obtained at day 0 (D0 baseline), 14 days post-vaccination 3 (D14 post-vaccine 3) and 5 or 6 days post-CHMI (D5/6 post-CHMI) (**Fig. 6A**). T cells were extracted into an individual Seurat object based on expression of canonical and lineage marker genes *CD3E*, *CD4*, *CD8A*, and *TRDC*. Clustering of the cells in this object revealed a total of 29 distinct clusters (**Fig. 6B**). Gene expression profiles were used to manually assign cell phenotypes to clusters (**Fig. S6A-B**). Given the observation that LARC GAP vaccination induces NKT-like γδ T cells in the murine liver and blood, we restricted our analyses to six clusters (8, 12, 18, 23, 24, and 26) containing candidate human γδ T cell/γδ NKT cell equivalents based on expression of *TRDC*, *TRGC1*, *TRGC2, FCGR3A* (encodes CD16), *NCAM1* (encodes CD56), and *KLRB1* (encodes CD161), all genes that have previously been identified in γδ T and NKT-like cells^41^ (**Fig. 6C-D, Fig. S6A-B**). We also analyzed a seventh cluster (cluster 2), which we classified as NK cells based on limited expression of *CD3E*.

**Figure 6:**
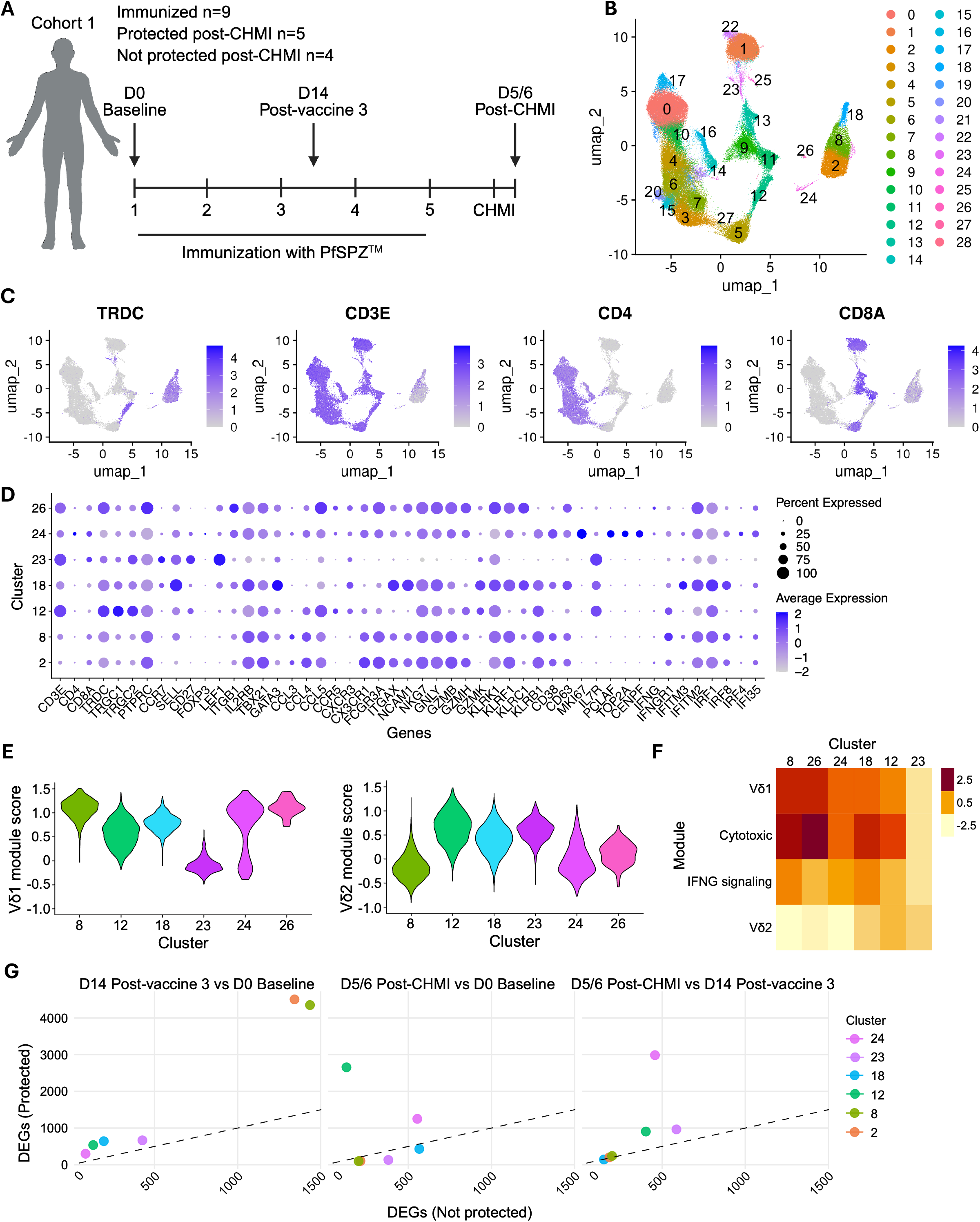
Human T and NKT-like cells in the blood following WPV immunization and CHMI. **A)** Immunization by Mosquito bite with RAS (IMRAS) trial volunteers received 5 PfSPZ^TM^ doses, followed by CHMI. Timepoints D0 baseline, D14 post-vaccine 3, and D5/6 post-CHMI indicate the days at which peripheral blood leukapheresis samples were taken from the nine study participants. **B)** Uniform manifold approximation and projection (UMAP) plot indicating clusters of distinct T cell subsets identified in the peripheral blood samples using single-cell RNA sequencing. Numbers indicate cell clusters. **C)** Expression levels of T cell markers on the UMAP plot of all identified cell clusters. **D)** Gene expression profiles for each of the γδ T/NKT and NK cell clusters. Dot color intensity indicates average expression level while dot sizes indicate percent of marker expression in the corresponding cluster. **E)** Violin plots showing aggregate gene module expression scores for Vδ1+ and Vδ2+ cells in each of the six clusters containing γδ T and NKT cells. **F)** Average Vδ1, Vδ2, cytotoxic/cytolytic, and IFNG-signaling gene module expression scores for each of the six clusters containing γδ T and NKT cells. See **Table S5** for a list of genes included in each module. **G)** Numbers of differentially expressed genes (DEGs) that significantly change (p value of <0.05) as a function of time and protection status in γδ T/NKT and NK cell clusters per timepoint comparison in protected and not protected individuals, calculated using mixed effect linear regression.

Given the preferential expansion of cytotoxic IFNγ+ γδ NKT cells in LARC GAP-immunized mice, we also assessed expression of chemokines including *CCL5*, cytotoxic/cytolytic markers including *GNLY*, *NKG7*, *KLRK1*, and *GZMB/H/K*, and tissue-homing markers *CXCR3*, *CCR5*, and *CX3CR1* (**Fig. 6D, Fig. S6A-B**) that have been shown to be expressed on liver-resident and liver-enriched γδ T cells in the human liver^42^. Since the 10x FLEX platform used to generate this data does not support detection of V(D)J sequences that would allow us to segregate these human γδ T/NKT cells into tissue-enriched Vδ1+/ Vδ3+ or blood-enriched Vδ2+ subsets, we leveraged published gene expression modules of human TCRVδ1 and TCRVδ2 γδ T cells^43^ to determine the relative identity of each cell cluster in our dataset (**Fig. 6E, Table S5**). We also generated gene expression modules that broadly report cytotoxic/cytolytic and IFNγ-signaling properties and measured their average module expression scores in each of the six clusters (**Fig. 6F, Fig. S6C, Table S5**). Together, these analyses revealed that two of the six clusters (8 and 26) expressed NK cell markers *FCGR3A* and *NCAM1*, were strongly cytotoxic/cytolytic, and contained cells with positive tissue-enriched Vδ1+ scores only (**Fig 6D-F**). Consistent with T cell tissue homing and patrol^44^, a high proportion of cells in each of these clusters also expressed *CX3CR1* (**Fig. 6D**). Of the two clusters, cluster 8 had the highest IFNγ-signaling score (**Fig. 6F**) and also expressed *KLRB1*, which has been associated with NKT cells that accumulate in the liver^45^ and with *Plasmodium*-responsive IFNγ-producing γδ T cells^46^. The four remaining clusters of γδ T and NKT-like cells (12, 18, 23, and 24) contained cells with positive Vδ2+ scores only or appeared to be mixed, containing cells with both positive Vδ1 and Vδ2 scores (**Fig. 6E-F**). Interestingly, cluster 24 showed both clearly defined Vδ1+ and Vδ2+ subpopulations and expressed a similar combination of NKT, cytotoxic/cytolytic, IFN-signaling, *CX3CR1*, and *KLRB1* markers as clusters 8 and 26. However, it was distinguished by cell cycle markers including *MKI67*, *PCLAF*, *TOP2A*, and *CENPF* (**Fig. 6D**). The frequencies of cells in each of the seven clusters characterized (i.e. the six γδ T/NKT cells and the NK cell cluster) in peripheral blood did not change significantly during immunization or after challenge in either protected or not protected vaccinees (**Fig. S7A**).

We next investigated transcriptomic responses to vaccination and CHMI in the seven cell clusters identified, in both protected and not protected IMRAS participants. Differentially expressed genes (**DEGs**) were identified by examining gene expression changes in each cluster over time in each group. Cluster 26 was excluded due to low cell numbers. We used mixed effect linear regression to account for within-individual variation with an FDR threshold of 0.05. The post-vaccine 3 timepoint was compared to the D0 baseline timepoint to determine changes induced by PfSPZ^TM^ vaccination, and the post-CHMI timepoint was compared to both the D0 baseline and post-vaccine 3 timepoints to determine changes induced by infection. All six remaining clusters had greater numbers of DEGs in protected individuals in response to vaccination, perhaps reflecting a stronger activation of NK and γδ T/NKT cells in protected individuals at the post-vaccine timepoint. This difference was particularly stark in clusters 2 and 8 (**Fig. 6G, Fig. S8**). NK cells have previously been shown to be associated with RAS-mediated protection^47–49^. Thus, greater activation of cluster 2 (identified as an NK cell cluster) in protected individuals is consistent with published studies and validates our bioinformatic approach. Cluster 8 shares the most similarities with IFNγ+/cytotoxic tissue-enriched γδ T/NKT cells (**Fig. 6E-F**). The genes with the greatest fold-change as a function of protection status between baseline and post-vaccination timepoints in clusters 2 and 8 revealed several DEGs associated with cell migration, and tissue infiltration in protected individuals (e.g., *CLTCL1*^50^, *MEP1B*^51^, *CD200R1*^52^, and *ST3GAL6*^53^ in cluster 2, and *DOCK8-AS1*^54^, *FBXL22*^55^, and *VIPR1*^56,57^ in cluster 8) (**Fig. S8**). In contrast, clusters 12, 23, and 24 had a higher number of DEGs in protected individuals following CHMI, when compared to the baseline and/or post-vaccination timepoints (**Fig. 6G, Fig. S8**).

To identify biological processes represented by gene changes observed following PfSPZ^TM^ vaccination and CHMI, we performed an independent GSEA^58^ on all gene expression changes for each cluster and timepoint comparison, using the previously published hallmark gene sets that comprise sets of coherently expressed genes in blood^59^ (**Fig. S7B**). Numerous gene sets were enriched in both protected and not protected individuals at each timepoint comparison, although more sets rose to our level of significance in not protected individuals in all comparisons. For example, cluster 8 significantly upregulated genes related to IFNγ response, oxidative phosphorylation, apoptosis, and allograft rejection post-vaccination in not protected vaccinees, but these trended down in this cluster in protected vaccinees. Together, our data indicate that cytotoxic/IFNγ-producing, tissue-associated γδ NKT cells respond to PfSPZ^TM^ vaccination in humans and do so differently in protected and not protected vaccinees.

### Activated Vδ2-cells correlate with protection in WPV-immunized humans

To extend our transcriptomic findings, we analyzed flow cytometric data for PBMCs from participants in both cohorts 1 and 2 of the IMRAS trial^40^ (**Fig. 7A**). We found no correlations between the frequencies of all γδ T cells, Vδ2+ cells, or Vδ2-cells, and protection (**Fig. S9A-C**). However, we observed positive correlations between increased frequencies of CD38+NKG2d+ (encoded by the *KLRK1* gene) γδ T cells, CD38+NKG2d+ Vδ2+ cells, and CD38+NKG2d+ Vδ2-cells and protection on days 3 and 7 following the third vaccination (**Fig. 7B-D**). Interestingly, while the frequencies of CD38+NKG2d+ γδ T cells and CD38+NKG2d+ Vδ2+ cells in protected individuals did not correlate with protection at timepoints after day 7 post-vaccine 3, the frequencies of CD38+NKG2d+ Vδ2-cells remained significantly elevated in protected vaccinees throughout the remainder of the immunization period, highlighting recently activated NKG2d-expressing Vδ2-cells as a bonafide correlate of WPV-engendered protection.

**Figure 7:**
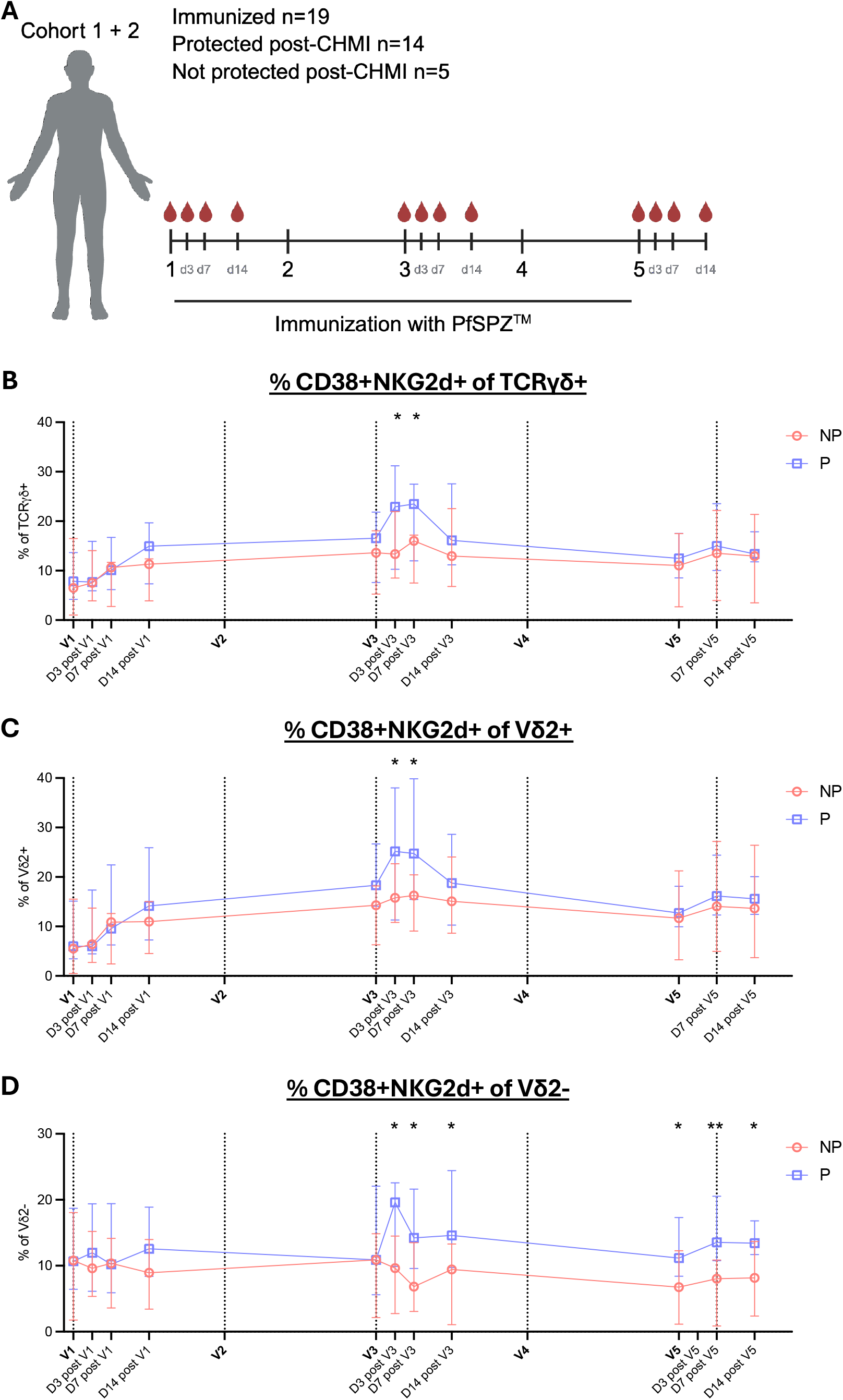
CD38+NKG2d+ Vδ2-cells correlate with WPV-engendered protection in humans. **A)** IMRAS trial volunteers received 5 vaccine doses. Peripheral blood mononuclear cell (**PBMC**) samples were taken from study participants at the timepoints indicated by the red blood droplets. **B-D)** Frequency of CD38+NKG2d+ cells in bulk TCRγδ+ cells (**B**), Vδ2+ cells (**C**), and Vδ2-cells (**D**). NP = not protected vaccinees, P = protected vaccinees. Plots show median ± 95% confidence intervals and data were compared at each timepoint using Welch’s t-test. *p<0.05, **p<0.01

## DISCUSSION

γδ T cells have been implicated in the protective immune response to promising anti-*Plasmodium* WPVs, but little is known about the subsets of γδ T cells induced by immunization or the distinct roles they play in the regulation of adaptive immunity to these WPVs. Late-arresting GAPs are significantly more efficacious than early-arresting WPVs in both preclinical and clinical studies^9–11^ and to date comprise the only malaria vaccine that can induce sterilizing immunity after a single immunization^11^. Thus, it is essential to examine the specific hepatic immune events that occur in response to LARC GAPs. Our study is the first to identify a role for hepatic γδ T cells in the induction of protective immune responses to LARC GAPs. We first examined the requirement for γδ T cells in the induction of sterile immunity by LARC GAPs and showed that in transgenic mice lacking γδ T cells, LARC GAP-engendered protective immunity was abrogated. This impaired protection was phenocopied in WT mice treated during immunization with a monoclonal antibody that causes the internalization of the γδ TCR^24^, but whether TCR recognition of parasite and/or host factors mediates γδ T cell activation and function during Plasmodium LS infection is unknown.

We next utilized single-cell mass cytometry to characterize the subsets of γδ T cells induced by LARC GAPs and showed that γδ T cells respond rapidly, heterogeneously, and across multiple anatomical sites to LARC GAP immunization, with the greatest changes occurring in the liver. A limit of our CyTOF approach is that our antibody panel did not contain the protein markers needed to conclusively identify γδ T cell-specific subsets. Therefore, although we observed 28 clusters of γδ T cells, many of these likely do not represent distinct functional subsets but rather different activation states, with the number of functional subsets likely to be much smaller. To develop a greater understanding of the γδ T cell subsets induced by LARC GAPs, we developed a γδ T cell-specific flow cytometry panel and observed that the number of hepatic γδ T cells doubled after immunization. This increase was primarily due to expansion of an IFNγ-expressing Vγ1+Vδ6.3+ population, which has been defined by others as γδ NKT cells^32,35^. These cells show phenotypic, functional, and transcriptional similarity to αβ NKT cells and are primarily localized in the liver and, to a lesser extent, the spleen^32,35,60^. To our knowledge, our study is the first to identify γδ NKT cells as the γδ T cell population preferentially induced in response to *Plasmodium* LS infection.

Given the requirement for γδ T cells in LARC GAP-induced protection and the robust induction of hepatic γδ NKT cells early after vaccination, we hypothesized that these cells may play a role in the regulation of the protective hepatic CD8+ T cell response. We found that functional inhibition of pan-γδ T cells or depletion of Vγ1+ cells alone abrogated the hepatic effector CD8+ T cell response to LARC GAP vaccination, highlighting a previously unappreciated functional role for Vγ1+ cells in the induction of protective hepatic T cells against LS parasites. Importantly, while hepatic CD8+ T cell responses to LARC GAP immunization are significantly impaired in the absence of these γδ NKT cells, splenic CD8+ T cells were not impacted. This is in stark contrast to a recent study where RAS immunization resulted in robust Vγ1+ cell activation in the spleen but muted and delayed kinetics of Vγ1+ cells in the liver and LDLN respectively, indicative of a role for their Vγ1+ cells in splenic priming of CD8+ T cells^22^. In comparison, we show robust γδ T cell activation in the liver with minor responses in the spleen and LDLN, supportive of a specific role in hepatic but not splenic CD8+ T cell responses. The disagreement between these two studies may result from differences in where naïve CD8+ T cells are primed in each system. Antigens generated during the LS account for the high protective efficacy of LARC GAP vaccines^9^. For bonafide LS antigens, antigen-presenting cells infiltrate the liver to acquire these antigens from *Plasmodium*-infected hepatocytes and then traffic to the LDLNs, not the spleen, to prime CD8+ T cells^5^. This is further supported by evidence from our group showing that splenic priming is dispensable for LARC GAP-engendered protection^13^. In contrast, this requirement for LDLN priming is not upheld for antigens that are also expressed on irradiated SPZs, such as the PbT-I/RPL6 or CS5M epitopes examined in the study by Le et al^22^. Further, high doses of irradiation, such as those used in Le et al.’s study, can result in reduced liver infiltration and/or invasion of hepatocytes, limiting antigen acquisition in the liver. Thus, splenic priming may have been more critical for CD8+ T cell responses in Le et al.’s model, while our use of LARC GAPs resulted in a situation in which hepatic CD8+ T cell responses were more impacted by γδ T cells than splenic CD8+ T cell responses. A careful analysis in both systems comparing the role of Vγ1+ cells in modulating T cell responses to SPZ antigens and LS-exclusive antigens may help in resolving these observed differences.

Nonetheless, both publications identify Vγ1+ cells as the subset responsible for driving CD8+ T cell responses to attenuated *Plasmodium* parasites. We further showed that Vγ1+Vδ6.3+ γδ NKT cells comprise the vast majority of Vγ1+ cells induced by LS parasites, implicating γδ NKT cells as the precise subset responsible for promoting the hepatic CD8+ T cell response to LARC GAP vaccination. We identified robust IFNγ production as a feature of the γδ NKT cells induced in our study, but γδ NKT cells can also produce IL-4 in addition to IFNγ^32^. IL-4 production was not examined here. IFNγ can positively regulate multiple stages of the adaptive CD8+ T cell response, including dendritic cell maturation, early expansion, and recruitment to the site of infection^61–63^ as well as T_RM_ differentiation and survival^64,65^. Whether IFNγ and/or IL-4 produced by γδ NKT cells modulates hepatic CD8+ T cell responses to LARC GAP vaccination remains to be examined.

We identified a liver-associated γδ T cell population as critical to LARC GAP-induced immunity. Obtaining liver samples from humans during clinical trials to detect correlates of these cells would be difficult. However, we made the striking observation that while IFNγ-producing γδ NKT cells are a minor population of γδ T cells in the blood, they are easily detectable after LARC GAP immunization in mice, supportive of a model in which activated γδ NKT cells traffic into circulation where they can be characterized as peripherally accessible mechanistic biomarkers of WPV-induced protection. There is no known human correlate of murine γδ NKT cells, and indeed direct human to murine correlation of γδ T cells is challenging given different receptor usage. To identify human cells with possible similarities to murine γδ NKT cells, we leveraged single-cell RNA sequencing to analyze lymphocytes from participants in cohort 1 of the IMRAS clinical trial. Out of 29 T lymphocyte clusters, we identified six clusters containing γδ T/NKT cells based on γδ TCR chain expression. Two of these clusters (8 and 26) exhibited increased expression of NK/NKT receptor, cytotoxic/cytolytic (including *KLRK1*), IFNγ-signaling, and tissue-homing transcripts, as well as an overall gene expression program that significantly overlaps with that of tissue-enriched human Vδ1+ γδ T cells. The other putative γδ T cell clusters (12, 18, 23, 24) exhibited Vδ2+ or a mixture of Vδ1+ and Vδ2+ programs, reflecting either a mixed population of γδ T cells, or Vδ2+ γδ T cells that share transcriptional signatures with Vδ1+ cells upon activation. Importantly, we observed that gene expression in cluster 8, the cell cluster that scored the highest in our Vδ1+, cytotoxic, and IFNγ modules, correlated with protection two weeks after the third PfSPZ^TM^ vaccination. To strengthen our novel observation of a positive correlation between Vδ1+ γδ T cell activation and protection, we leveraged high-parameter flow cytometry to analyze Vδ2+ and Vδ2-γδ T cell populations in PBMCs collected from both cohorts 1 and 2 of the IMRAS trial. Critically, we found that Vδ2-cells co-expressing the T cell activation marker, CD38^66^, and the NK cell receptor, NKG2d (encoded by *KLRK1*), positively correlated with protection at every timepoint tested following the third immunization. In contrast, a positive correlation of CD38+NKG2d+ cells in the Vδ2+ population was only observed on days 3 and 7 post vaccination dose 3, indicating that Vδ2-γδ T cell populations might be a more robust biomarker of WPV-engendered protection. NKG2d is expressed on cytotoxic γδ T cells, is utilized in stress ligand sensing, and is expressed on human IFNγ-producing γδ T cells that respond to *Plasmodium* blood-stage parasites^67,68^. Whether the presence of these markers on the Vδ2-cells observed here marks a unique functional subset or simply a state of activation is unclear. Together our analyses of human PBMCs demonstrate that subsets of γδ T cells bearing characteristics of IFNγ+ tissue-associated γδ NKT cells are responsive to early-arresting WPVs in humans and correlate with protection.

It is important to note that the role of γδ T cells in WPV-induced protection in humans is likely complex and will require further analyses to fully characterize the human correlates of the γδ NKT cells identified in our mouse model. Mouse γδ NKT cells express the Vγ1+Vδ6.3+ TCR, a semi-invariant TCR that is conceptually similar to the semi-invariant human Vγ9+Vδ2+ TCR. However, murine γδ T cells are not known to recognize phosphoantigens like Vγ9+Vδ2+ cells^69^ and while Vγ9+Vδ2+ cells are primarily found in the periphery^23^, murine γδ NKT cells are tissue-enriched. Human Vδ1+ cells are primarily found within tissues^23,42^ and are abundant in the blood of individuals from malaria-endemic regions and those who have undergone repeated blood-stage *P. falciparum* infection^70–72^. However, Vδ1+ cells have more variable TCRs and are considered to be more adaptive-like^73^. Therefore, further characterization of murine γδ NKT cells and human γδ T cell subsets, especially within LARC GAP-immunized hosts, is needed.

Our study is the first to identify murine hepatic γδ NKT cells as the subset that primarily responds to and regulates the protective hepatic CD8+ T cell response to highly protective LARC GAPs. Analyses of WPV-immunized individuals also identified both human Vδ2+ and Vδ2-γδ T cells as responsive to WPVs. We show for the first time that activation of Vδ2-γδ T cells positively correlates with protection, supporting a rationale for characterizing these tissue-associated γδ T cells as biomarkers of protective immunity in future clinical trials, especially within malaria-endemic populations. Additionally, a better understanding of how these γδ NKT cells and their human correlates are induced, what they respond to, and precisely how they influence conventional T cell responses in the liver might be useful in the design of γδ T cell-targeted adjuvants that enhance hepatic CD8+ T cell responses.

## MATERIALS/METHODS

### Ethics statement

Animal studies were performed according to the regulations of the institutional animal care and use committee (IACUC). Approval was obtained from the Seattle Children’s Research Institute IACUC under protocol 00697. The Seattle Children’s Research Institute IACUC adheres to the NIH Office of Laboratory Animal Welfare standards (OLAW Public Health Service Assurance #D16-00119).

For the IMRAS trial, written informed consent was obtained from all study participants. Ethics approval was obtained from the Naval Medical Research Center (NMRC) Institutional Review Board. The study was conducted according to the Declaration Helsinki and Good Clinical Practices under the guidelines of United States Food and Drug Administration Investigational New Drug (IND) application BB-15767. The trial was registered on ClinicalTrials.gov (NCT01994525). Details of the study design for the IMRAS trial have been described previously^40^.

### Mice

Female Swiss Webster mice for parasite cycles were purchased from Envigo laboratories. Six-to 8-week-old female C57BL/6 (**B6,** 000664), GREAT (017581), and TCRδ^−/-^ (002120) mice were purchased from Jackson Laboratories. TCRδ-eGFP mice were kindly provided by Dr. Karen Edelblum. Mice were maintained and bred under specific pathogen-free conditions at Seattle Children’s Research Institute. Mice were euthanized by CO_2_ asphyxiation and cervical dislocation prior to organ isolation or at the end of each experiment.

### Sporozoite isolation

Mosquitos were reared through the ACL-1 insectary at Seattle Children’s Research Institute. 8-to 16-week-old female Swiss Webster mice were injected intraperitoneally (**i.p.**) with blood-stage *Plasmodium yoelii* (***Py***) *luc*^26^ or *Py fabb/f^−^*^25^ (LARC GAP). Three days later, gametocyte exflagellation was confirmed and the infected mice were used to feed female *Anopheles stephensi* mosquitos. 14 to 16 days after the blood meal, salivary glands harboring sporozoites (**SPZs**) were isolated from the mosquitos and used in mouse infections. For mock infections, salivary glands from uninfected mosquitos were prepared in the same manner.

### Mouse infections

For adaptive vaccine studies, 8-to 16-week-old female B6 and TCRδ^−/-^ mice were immunized intravenously (**i.v.**) via retro-orbital plexus injection with 50,000 LARC GAP SPZs or uninfected mosquito salivary glands. This immunization was repeated 28-49 days later. Immunized mice were then challenged 38-42 days after the secondary immunization with 10,000 *Py luc* SPZs. Parasitemia was monitored by Giemsa-stained thin smears beginning on day 3 post-challenge. At least 20 fields of >200 red blood cells per field were examined for patency determination. Mice showing no evidence of blood stage infection by day 16 post-infection were assessed as sterilely protected. For mass cytometry and flow cytometry experiments examining γδ T cells, 8-to 16-week-old female B6 or GREAT mice were immunized i.v. with 100,000 LARC GAP SPZs or uninfected mosquito salivary gland debris as a control. For examination of CD8+ T cell responses post-vaccination, 8-to 16-week-old female B6 mice were immunized i.v. with 50,000 LARC GAP SPZs or uninfected mosquito salivary glands.

### Murine organ harvests, cell isolation, and flow cytometry

#### Liver

Liver lobes were perfused through the hepatic vein using HBSS with HEPES followed by a 0.5 mg/mL collagenase solution. Perfused liver lobes were manually dissociated through 100 µM nylon filters using the plunger of a 10 mL syringe into PBS with 4% fetal bovine serum (**FBS**). Cells were spun at low speed to remove hepatocytes. Non-parenchymal cells were isolated using a 40% iodixanol gradient and spun at 1500 rpm for 25 minutes with no brake. Cells were collected in PBS + 2% FBS for downstream antibody staining.

#### Blood

Heparinized blood was collected via cardiac stick to the heart and treated with ACK lysis buffer to remove red blood cells. Cells were washed with PBS + 2% FBS and collected in the same buffer for downstream antibody staining.

#### Lymph Nodes

Liver-draining lymph nodes (**LDLNs**) were manually dissociated using the plunger of a 1 mL syringe into PBS + 2% FBS and the cells were collected for downstream antibody staining.

#### Spleen

Spleens were manually dissociated through 100 µM nylon filters using the plunger of a 10 mL syringe into PBS with 2% FBS and washed. The cell pellets were incubated with ACK lysis buffer to remove red blood cells followed by a final wash. Cells were collected in PBS + 2% FBS for downstream antibody staining.

#### Antibody staining & flow cytometry

Isolated cells were incubated with Biolegend mouse FcX block (catalog no. 101320) for 15 minutes at 4°C to block non-specific staining. Cells were then stained with surface antibodies at 4°C for 30-45 minutes (see **Table S3**). When performing intracellular staining or for analysis of CD8+ T cells, cells were fixed/permeabilized using the BD Cytofix/Cytoperm Fixation/Permeabilization Kit (catalog no. 554714) per the manufacturer’s instructions. For intracellular staining, cells were stained with intracellular antibodies at 4°C for 60 minutes (see **Table S3**). Labeled cells were run on a BD FACSymphony A5 cytometer (RRID:SCR_022674) and analyzed with FlowJo v10.10 software (BD Life Sciences). All data were reported as the number of cells per 1 million live singlet lymphocytes.

### Mass cytometry and data processing

Perfused livers, LDLNs, and spleens were harvested from B6 mice infected with LARC GAPs (n=5) or uninfected mosquito salivary glands (**mock**) (n=5) at 4 hours or on days 3, 7, and 14 post-immunization, and dissociated to isolate non-parenchymal cells as described above. Prior to staining, the samples were barcoded using Cell-ID 20-Plex Pd barcode kit (catalog no. 201060, Standard Biotools, CA) as per manufacturer’s protocol. These samples were then stained with metal conjugated antibodies (**Table S4**) against various lineage and CD8+ T cell markers and run on the Helios CyTOF systems (Fluidigm). Metal conjugation for all markers except CD45 (see **Table S4** for details) was performed as described previously^74^ using either the MAXPAR DN3 (catalog no. 201144C, Standard Biotools, CA) or the MCP9 (catalog no. 201116A, Standard Biotools, CA) antibody labeling kit. Cisplatin (platinum isotopes 194/195) (catalog no. 201064, Standard Biotools, CA) was used to stain dead cells before fixation. Cell were fixed using 2% PFA overnight at 4°C, and frozen at -80°C. Cells were then retrieved in MAXPAR cell acquisition buffer (catalog no. 201240, Standard Biotools, CA) and immediately acquired by mass cytometry with an acquisition rate of 200-250 cells/sec. 2% of EQ Four Element Calibration Beads (catalog no. 201078, Standard Biotools, CA) were added into the cell suspension in order to normalize the signal variation. Bead normalization was performed on raw data using premessa tool in R. Data processing was performed using FlowCore (R), which included compensating for spillover and batch correction by transforming data (arcsinh transformation) to adjust for signal distribution. Next, quality control was performed by filtering out doublets, dead cells, and low-quality events to ensure accurate downstream analysis. The processed data was then subjected to clustering and population identification using FlowSOM (R), to help define distinct immune or cellular subsets based on marker expression. Differential abundance and functional analysis were conducted to compare population frequencies and assess activation states. To interpret the results, phenographs, heatmaps, and dot plots were created to provide insights into marker expression patterns and immune landscape profiling across samples. When statistically comparing cluster frequency in mock-vs. LARC GAP-immunized mice, significant differences were only reported if average cluster frequency was at least 1%.

### Antibody treatment

For inhibition of γδ T cells in adaptive vaccine studies, 0.2 mg GL3 (BioXCell, contract production) was injected i.p. into mice at days -1 and 27. For inhibition of γδ T cells and depletion of Vγ1+ cells when studying CD8+ T cell responses, 0.2 mg GL3, α-Vγ1 (BioXCell, BE0257), or Armenian Hamster IgG (BioXCell, BE0091) were injected i.p. into mice in the morning prior to vaccination. To confirm depletion of Vγ1+ cells, 0.2 mg α-Vγ1 was injected i.p. into mice 24 hours prior to organ harvest.

### *In vivo* bioluminescent imaging

Mice infected with *Py luc* were imaged at 40-44 hours post-infection. Each mouse was injected with 2.25 mg luciferin (Gold Biotechnologies, LUCK-1G) diluted in PBS intraperitoneally. Following a 5-minute anesthetized incubation period (XGI-8, Caliper Life Sciences), mice were placed in the Perkin Elmer *in vivo* imaging system (**IVIS**) (RRID:SCR_018621) and imaged for 3 minutes. Total flux values for each mouse were determined using Living Image analysis software (Perkin Elmer).

### Human PBMC flow cytometry analysis

Flow cytometry profiling, panels, and gating strategies for these samples have been described previously^49,75,76^. Briefly, PBMCs were collected from trial participants and stored frozen prior to thawing for analysis in RMPI with 10% FBS and benzonase nuclease. Cells were incubated with LIVE/DEAD Fixable Blue Dead Cell Stain Kit and Human BD Fc Block for 30 minutes at room temperature and then stained with previously described flow panels^49,75,76^. Labeled cells were run on a BD FACSymphony flow cytometer and analyzed with FlowJo v10.10 software (BD Life Sciences). Live singlet cells were gated on prior to further gating, and all data were reported as frequencies.

### Single-cell RNA sequencing, data processing, and bioinformatics analyses

Peripheral blood leukapheresis samples from vaccinated individuals (n = 9) in cohort 1 of the IMRAS trial^40^ who were protected (n = 5) or not protected (n = 4) from CHMI, obtained at day 0 (D0 baseline), 14 days post-vaccination 3 (D14 post-vaccine 3) and 5 or 6 days post-CHMI (D5/6 post-CHMI). 10x Genomics Chromium Fixed RNA profiling was used for single-cell RNA-sequencing of these samples with a healthy human PBMC (BioIVT) bridging control included for cross-batch normalization downstream. In brief, up to one million cells per sample were processed and fixed, and probes were hybridized per the manufacturer’s instructions. Samples were then pooled into three batches, with the bridging control, at equivalent cell concentrations, and the single cell suspension pools loaded across three wells of Chip Q for gel beads-in-emulsion generation, and subsequent library generation. Cells were loaded at a concentration of 25,000 cells per barcoded sample. Libraries were sequenced using a NovaSeq X 10B flow cell, at a target depth of 10,000 reads per cell at Northwest Genomics Center at the University of Washington. Samples were then computationally resolved and quality-checked using in-house pipelines.

The binary base call (BCL) sequence files were base-called and demultiplexed using cellranger mkfastq. GEM wells for each sample were aggregated and depth normalized using cellranger aggr. A Seurat (v5) object was created for each sample following debris removal and barcode mapping to samples. Raw count data was pre-processed to remove cell doublets (via Scrublet^77^) and low quality cells with high mitochondrial RNA fraction (10%) and very low or very high gene counts (nFeature_RNA <200 and >5000). The per sample RNA data were independently normalized using standard normalization, then each integrated across all samples using the Harmony method^78^ to correct for any batch effects and control for between-sample technical variation. Manual annotation of cellular identity was performed by finding differentially expressed genes for each cluster using Seurat’s implementation of the Wilcoxon rank-sum test (“FindMarkers” function) and comparing those markers with known cell type–specific genes^43^. The T and NK cell clusters were then extracted into individual Seurat objects and analyzed as above. After all filtering and cluster extraction steps, the dataset reported here contains a total of 149,816 cells, with an average of ∼16,646 cells (minimum 9,344, maximum 22,477) per sample.

To calculate differentially expressed genes (DEGs) in individual clusters between timepoints in protected and not protected individuals, the raw gene expression counts per cluster were aggregated to generate pseudobulk count matrices. A linear mixed-effect regression model (LMER) was then fit for each cluster through the R package glmmSeq^79^. Gene expression (counts per million) was fit as a dependent variable while sample timepoint and protection status were fit as fixed effects, and individual identifier was a random effect. Gene set enrichment analysis (GSEA) was performed using the R package fgsea^80^. The MSigDB hallmark gene sets^59^ were obtained from download site http://www.gsea-msigdb.org/gsea/msigdb/genesets.jsp?collection=H. Differentially expressed genes for each comparison were ranked by their signed P values across different time interval comparisons and separately for each sample groups. Gene sets with normalized enrichment scores (NES) between -1 and 1 were adjusted to 0. Statistically significant gene sets were identified as those with less than 0.05 False Discovery Rate (FDR). FDR-corrected P values were calculated using the Benjamini-Hochberg method.

### Statistical analyses

Analyses of murine data and human flow cytometry data were performed using GraphPad Prism Software (Prism 10 for macOS, Boston, MA). For comparison of multiple groups, statistical significance was determined using non-paired one-way Brown-Forsythe and Welch ANOVA tests followed by Dunnett’s T3 multiple comparisons test. Kaplan–Meyer patency curves were compared via the log rank Mantel–Cox test. Outliers were identified in IVIS results utilizing Grubb’s test. All other statistical significance was determined using the non-paired two-tailed Welch’s t-test.

## DATA AVAILABILITY

Processed scRNA-seq data is available on Figshare 10.6084/m9.figshare.31712581.

## ACKNOWLEDGEMENTS

We thank members of the Minkah Lab, the McDermott Lab, the Newell Lab, and the Allen Institute for Immunology for their technical assistance and helpful comments. We would like to acknowledge all the IMRAS study participants who made this work possible. We thank the Naval Medical Research Center (NMRC) Malaria Department, the Clinical Trials Center, and the Clinical Immunology Laboratory staff led by Dr. Eileen Villasante, Dr. Judith Epstein, Dr. Martha Sedegah, respectively, that conducted the IMRAS clinical trial and processed, cryopreserved, inventoried, and transferred the clinical samples. This work was supported by institutional funds awarded to Dr. Minkah and Drs. Stuart, Newell, Skene, and McDermott, 5R01AI170777, and 5U19AI128914 from the National Institutes of Health.

## AUTHOR CONTRIBUTIONS

Conceptualization: RCB, AVK, SMM, NKM. Data curation: RCB, AVK, NH, LG. Formal analysis: RCB, AVK, NH, CP, LG, SMM. Investigation: RCB, AVK, AAK, GJ, SMN, NS, EGT, NH, KVS, SCDR, CP, LG, SMM. Software: AVK, AND, LG. Visualization: RCB, AVK, AND, CP, LG, SMM. Statistical analyses of clinical trial data: RCB, CP, SMM. Access to all clinical trial data and samples: NH, SCDR, KDS, EWN, SMM. Access to all clinical trial data generated and analyzed: KDS, SMM. Writing – original draft: RCB, SMM, NKM. Writing – review & editing: RCB, AVK, AND, AAK, GJ, SMN, KDS, SMM, NKM. Supervision: EWN, SMM, NKM. Funding acquisition: KDS, SCDR, PJS, EWN, SMM, NKM. All authors agreed to manuscript submission, read and approved the final draft, and take full responsibility of the content, including data accuracy and statistical analyses.

## DECLARATIONS OF INTEREST

The authors declare no competing interests.

---

**Figure S1: Expression of protein and intracellular markers on hepatic γδ T cells at day 4 post-immunization**

Mice were immunized with LARC GAP SPZ or mock and livers were harvested for flow cytometry analysis at day 4 post-immunization. **A)** CD62L+ γδ T cells per 1 million live singlet lymphocytes in the liver. **B)** CD69+ γδ T cells per 1 million live singlet lymphocytes in the liver. **C)** CD122+ γδ T cells per 1 million live singlet lymphocytes in the liver. **D)** CD160+ γδ T cells per 1 million live singlet lymphocytes in the liver. **E)** ICOS+ γδ T cells per 1 million live singlet lymphocytes in the liver. **F)** Ki67+ γδ T cells per 1 million live singlet lymphocytes in the liver. **G)** IFNγ+ γδ T cells per 1 million live singlet lymphocytes in the liver. All dot plots show the median ± 95% confidence interval, and each dot represents one mouse. Data were compared using Welch’s t-test. Data for **A-E** come from at least two independent experimental replicates and data from **F, G** are from one replicate. *p<0.05, **p<0.01, ****p<0.0001

**Figure S2: Vγ1+Vδ6.3-cells in the liver and blood at day 4 post-immunization**

Mice were immunized with LARC GAP SPZ or mock and organs were harvested for flow cytometry analysis at day 4 post-immunization. **A)** Vγ1+Vδ6.3-cells per 1 million live singlet lymphocytes in the liver. **B)** Number of YFP+ and YFP-Vγ1+Vδ6.3-cells per 1 million live singlet lymphocytes in the liver. **C)** Vγ1+Vδ6.3-cells per 1 million live singlet lymphocytes in the blood. **D)** Number of YFP+ and YFP-Vγ1+Vδ6.3-cells per 1 million live singlet lymphocytes in the blood. All dot plots show the median ± 95% confidence interval, and each dot represents one mouse. Red numbers above each plot show the fold-change difference between the selected groups. For stacked bar plots showing YFP+ vs. YFP-cells, median values ± 95% confidence intervals are shown. Data were compared using Welch’s t-test. All data come from at least two independent experimental replicates. ****p<0.0001

**Figure S3: IFNγ expression in Vγ4+ and Vγ6+ cells in the liver at day 4 post-immunization**

Mice were immunized with LARC GAP SPZ or mock and livers were harvested for flow cytometry analysis at day 4 post-immunization. **A)** Percent of liver Vγ4+ cells that are YFP+ or YFP-. **B)** Percent of liver Vγ6+ cells that are YFP+ or YFP-. Normalized median values ± 95% confidence intervals are shown. All data come from at least two independent experimental replicates.

**Figure S4: CD4+ T cell responses in immunized γδ T cell-deficient mice**

Mice were treated with an IgG control antibody, ɑ-TCRγδ (**A,B**), or ɑ-Vγ1 (**C,D**) antibody on day 0 prior to mock or LARC GAP immunization. 7 days later, organs were harvested for flow cytometry. **A, C)** Number of CD4+ T cells per 1 million live singlet lymphocytes in the liver and spleen. **B, D)** Number of CD4+CD44+ T cells per 1 million live singlet lymphocytes in the liver and spleen. All plots show the median ± 95% confidence interval. Each dot represents one mouse. The Brown-Forsythe and Welch ANOVA test followed by Dunnett’s T3 multiple comparisons test were used to statistically analyze the data. All data come from at least two independent experimental replicates. ***p<0.001

**Figure S5: Depletion of Vγ1+ cells leads to a reduction in CD3+TCRβ-cells**

TCRδ-eGFP mice were treated with IgG or anti-Vγ1 one day prior to harvest of livers for flow cytometry analysis. Plot shows number of eGFP+CD3+TCRβ-cells per 1 million live singlet lymphocytes in the liver.

**Figure S6: Distinct T and NKT cell subset identities in WPV-immunized human PBMC samples were generated based on their gene expression profiles.**

**A)** Distinct T, NKT, and NK cell cluster identities in WPV-immunized human peripheral blood leukapheresis samples were generated using the Seurat package based on their gene expression (RNA) profiles. The dot plot represents the gene expression profiles for each of the clusters. Dot color intensity indicates average expression level while dot sizes indicate percent of marker expression in the corresponding cluster. **B)** Expression levels of specified marker genes on the UMAP plot of all identified cell clusters. **C)** Expression levels of Vδ1, Vδ2, cytotoxic/cytolytic, and IFNG-signaling gene modules on the UMAP plot of all identified cell clusters.

**Figure S7: Cell abundance and gene expression changes in human γδ T and NKT cells in the blood following WPV immunization and CHMI**

**A)** Percentage of cells of total T cells per cluster and per timepoint in protected vs. not protected participants. Dots denote means and error bars represent one standard deviation. **B)** Heatmap plot showing GSEA on comparisons of DEGs between indicated timepoints, individually tested in each of the γδ T/NKT and NK cell clusters from the protected and not protected IMRAS trial participants. Each square indicates a gene set. Color represents gene sets that are changing with normalized enrichment score (NES) < -1 and > 1 obtained from GSEA. A white dot in a colored square indicates gene sets with statistically significant changes. Grey represents gene sets that do not change, and white indicates gene sets that could not be quantified in each cluster. All gene expression changes were used as inputs for GSEA. Threshold for significant gene sets was set to an FDR-adjusted P value of <0.05.

**Figure S8: Gene expression changes in human γδ T and NKT cells in the blood following WPV immunization**

Numbers of differentially expressed genes (DEGs) that significantly change as a function of time (cyan), and time and protection status (ochre and magenta) in NK and γδ T cell clusters per timepoint comparison in protected and not protected individuals, calculated using mixed effect linear regression. The top 10 genes with the greatest fold-change as a function of time and protection status are labeled for each of the protected and not protected groups.

**Figure S9: Frequency of TCRγδ+ cells, Vδ2+ cells, and Vδ2-cells in IMRAS participants A-C)**

Frequency of TCRγδ+ cells (**A**), Vδ2+ cells (**B**), and Vδ2-cells (**C**) in CD3+ cells from IMRAS participants. NP = not protected vaccinees, P = protected vaccinees. Plots show median ± 95% confidence intervals.

**Figure S10: Total numbers of *Plasmodium*-induced γδ T cells in the liver four days post-immunization**

Mice were immunized with LARC GAP SPZ or a mock immunization and livers were harvested for flow cytometry analysis at day 4 post-immunization. **A)** Total γδ T cells in the liver. **B)** Total Vγ1+Vδ6.3+ cells in the liver. **C)** Total YFP+ γδ T cells in the liver. **D)** Total YFP+ Vγ1+Vδ6.3+ cells in the liver. All dot plots show the median ± 95% confidence interval, and each dot represents one mouse. Data were compared using Welch’s t-test. All data come from at least two independent experimental replicates. **p<0.01, ***p<0.001

**Figure S11: Flow cytometry gating strategies**

**A)** Gating strategy for murine γδ T cells, using representative flow cytometry plots from liver cells extracted from LARC GAP-immunized GREAT mice at day 4 post-immunization. **B)** Gating strategy for murine CD4+ and CD8+ T cells, using representative flow cytometry plots from liver cells extracted from LARC GAP-immunized B6 mice at day 7 post-immunization. **C)** Gating strategy for human γδ T cells, using representative flow cytometry plots from a not protected vaccinee at day 14 post-vaccine 3.

## Notes

### Competing Interest Statement

The authors have declared no competing interest.

