## Supplemental Figures for "Hepatic γδ NKT cells modulate liver-resident CD8+ T cells to attenuated malaria parasite vaccines"

#### Figure S1: Expression of protein and intracellular markers on hepatic γδ T cells at day 4 post-immunization

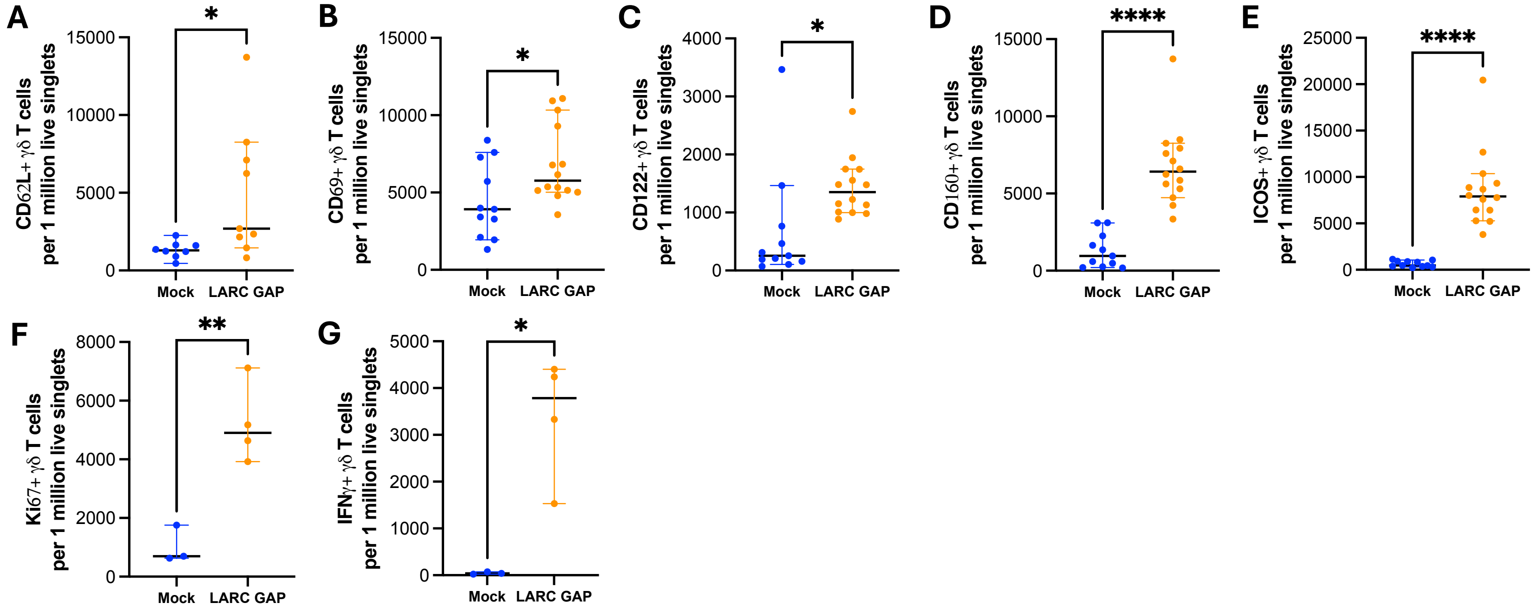

**Figure S2: Vγ1+Vδ6.3- cells in the liver and blood at day 4 post-immunization**

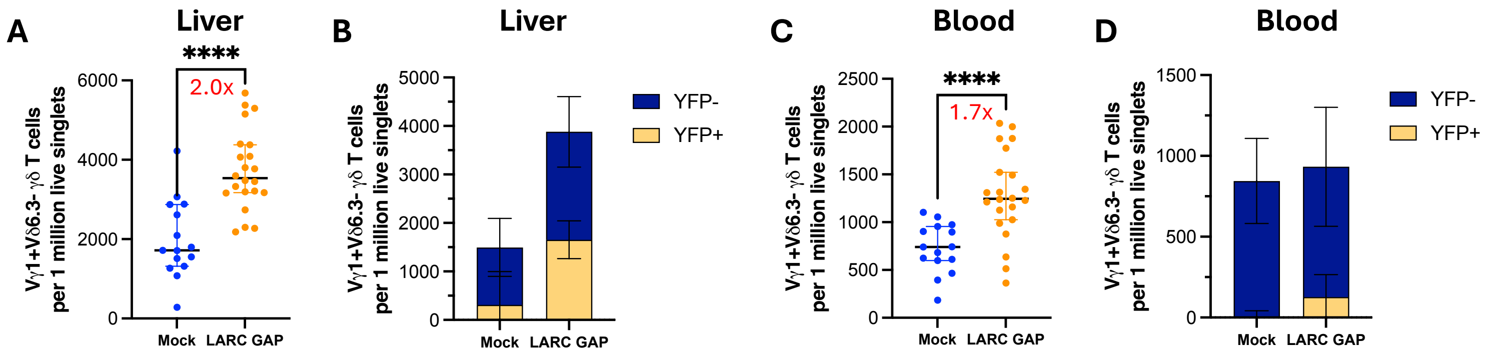

#### Figure S3: IFNγ expression in Vγ4+ and Vγ6+ cells in the liver at day 4 post-immunization

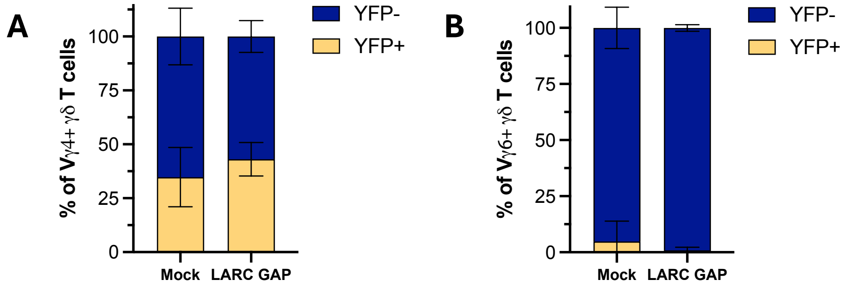

**Figure S4: CD4+ T cell responses in immunized γδ T cell-deficient mice**

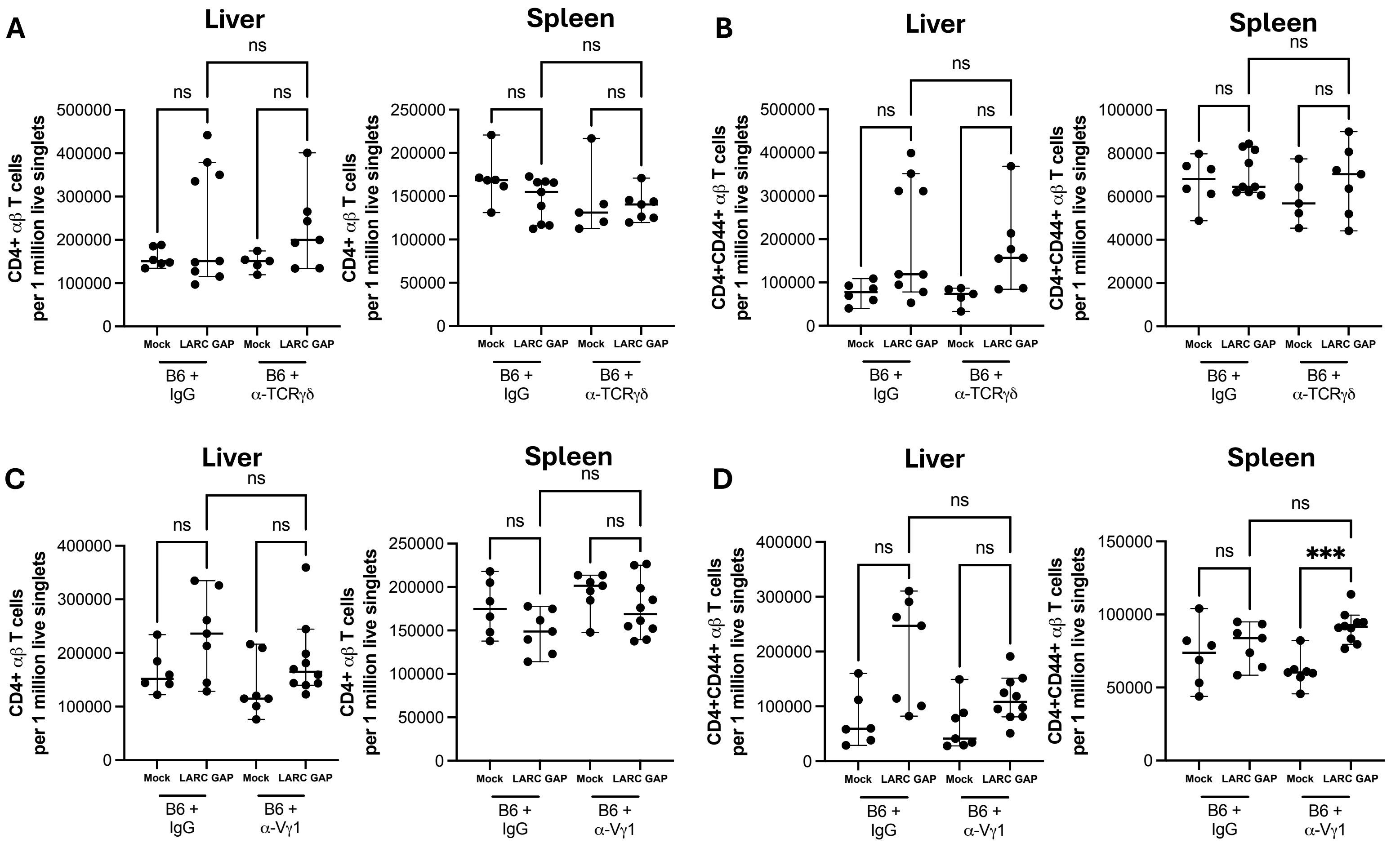

#### Figure S5: Depletion of Vγ1+ cells leads to a reduction in CD3+TCRβ- cells

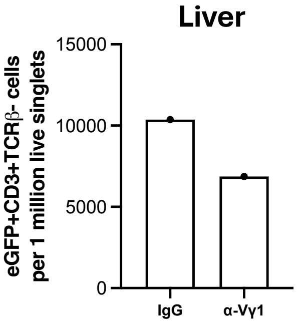

**Figure S6: Distinct T and NK cell subset identities in WPV-immunized human PBMC samples were generated based on their gene expression profiles.**

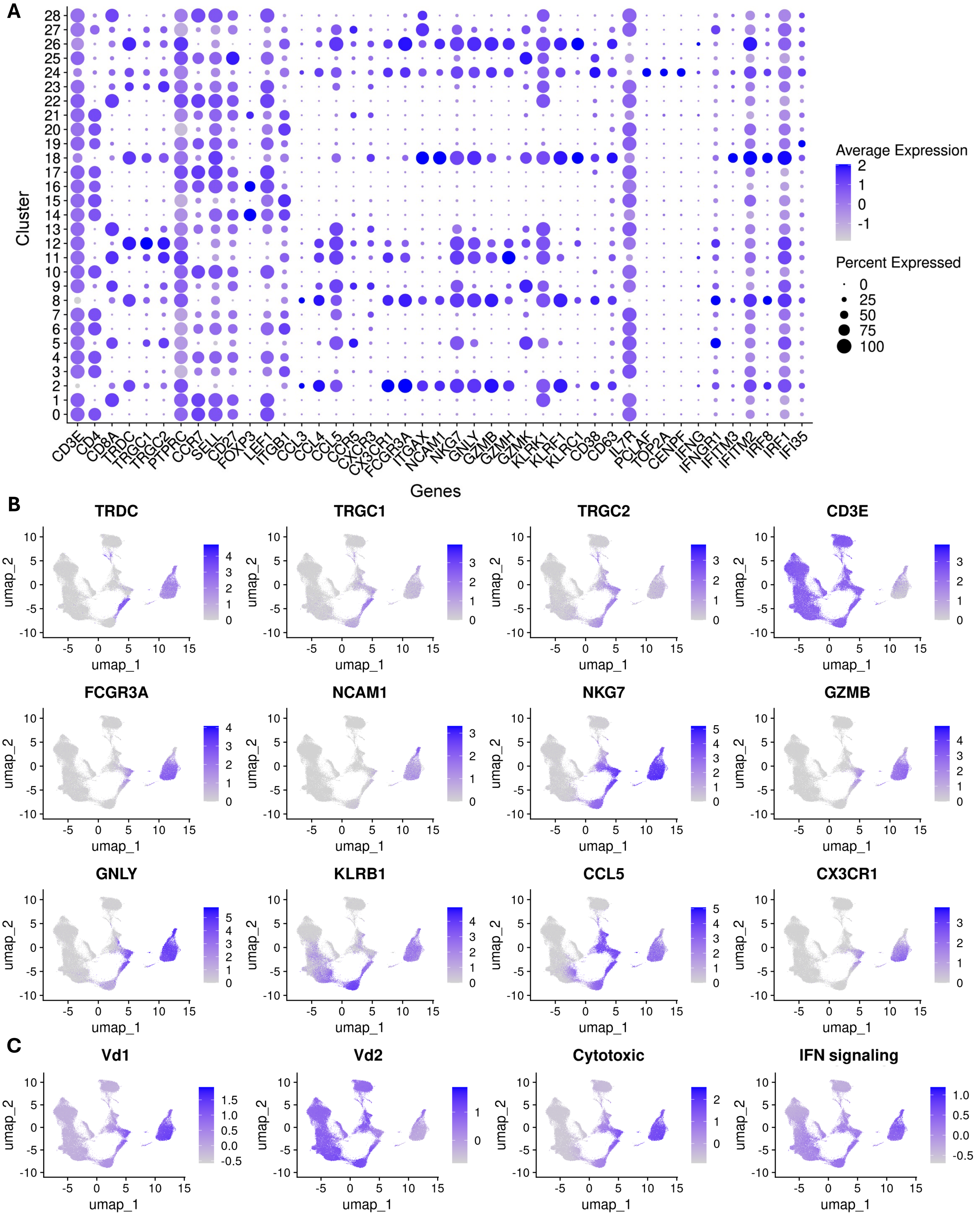

#### Figure S7: Cell abundance and gene expression changes in human γδ T and NKT cells in the blood following WPV immunization and CHMI

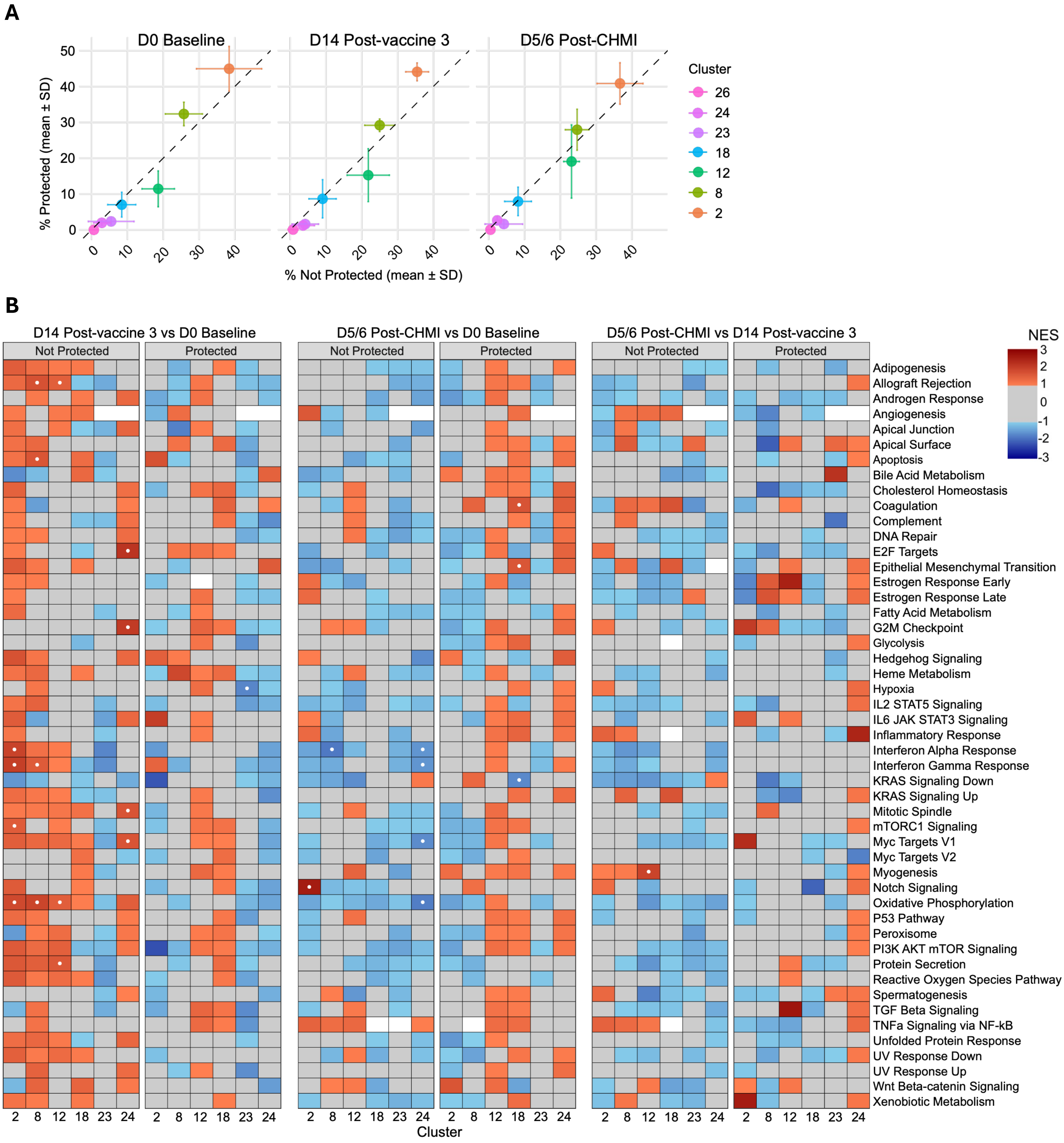

#### Figure S8: Gene expression changes in human γδ T and NKT cells in the blood following WPV immunization

**Cluster 2** D14 Post-vaccine 3 vs D0 Baseline

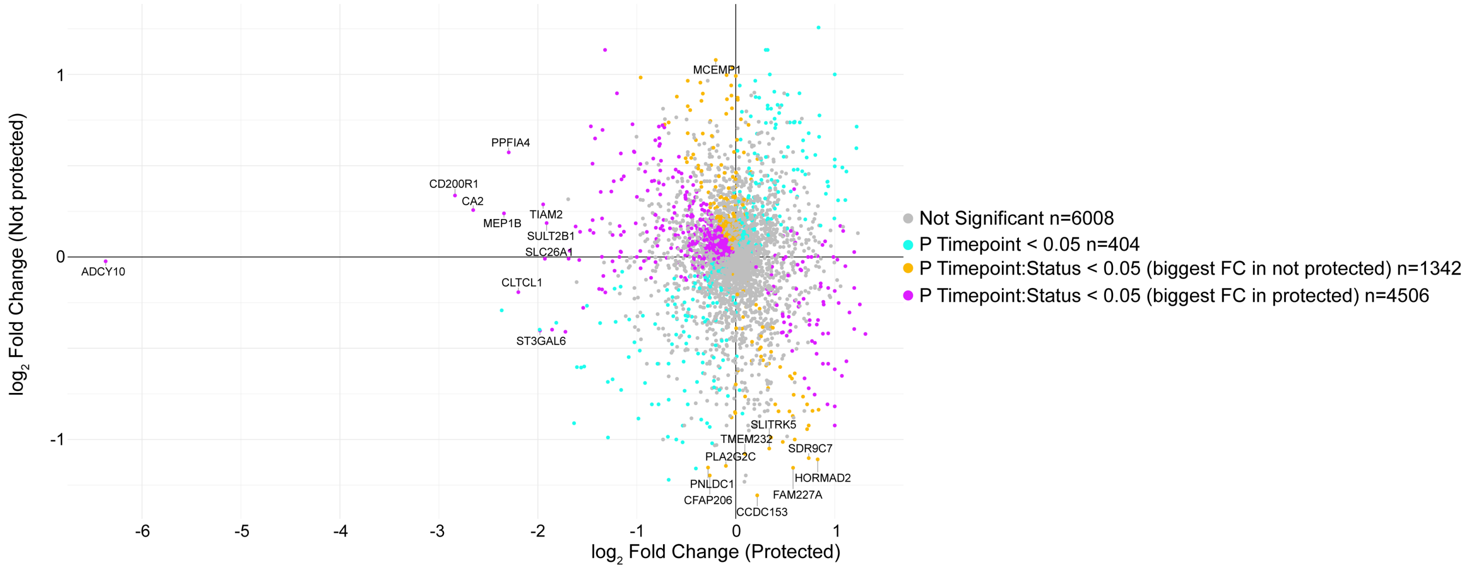


**Cluster 2** D5/6 Post-CHMI vs D0 Baseline

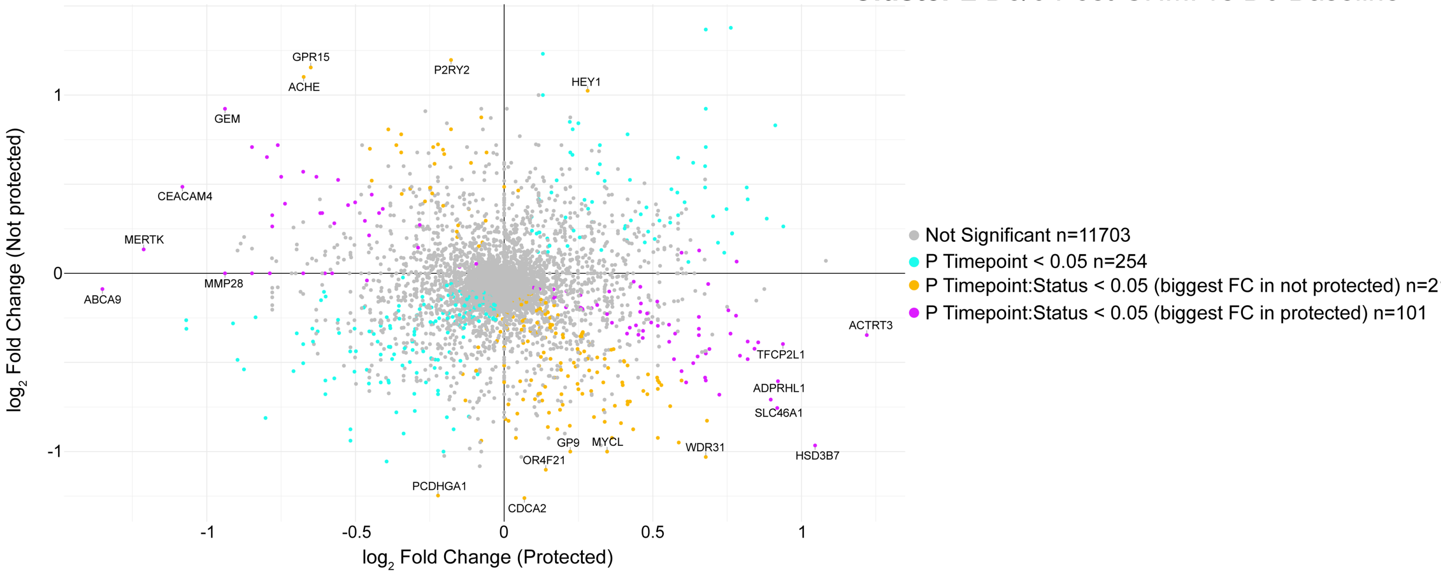


**Cluster 2** D5/6 Post-CHMI vs D14 Post-vaccine 3

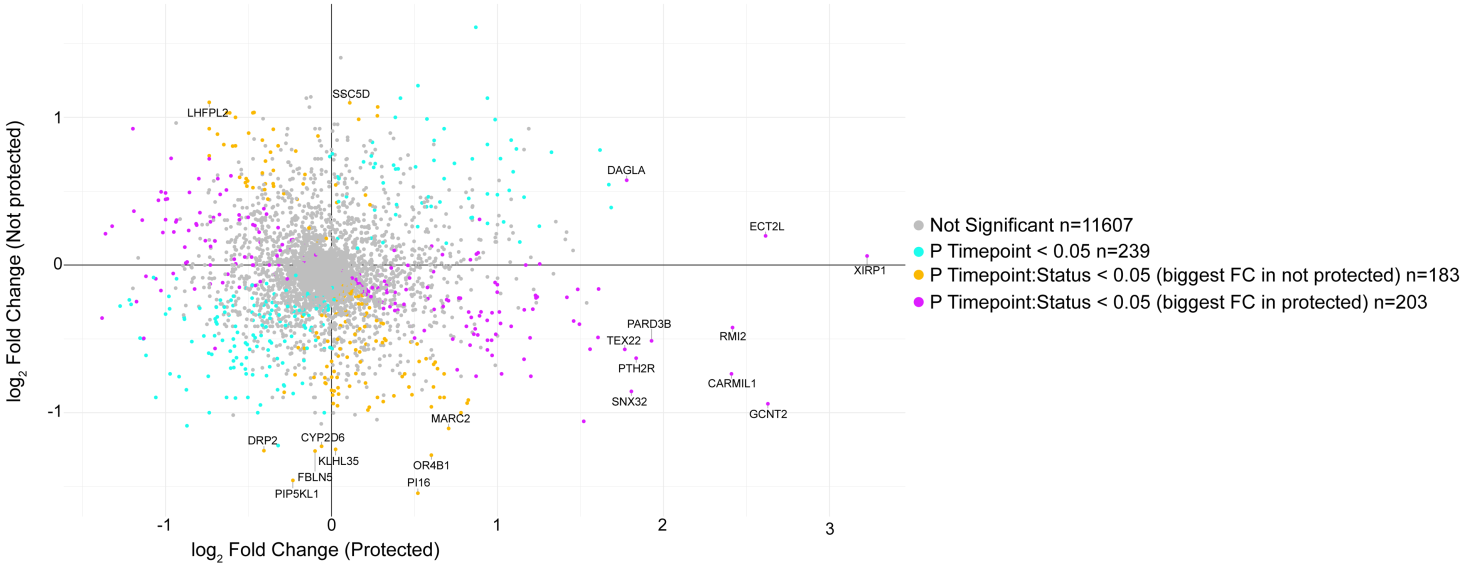

#### Figure S8 (continued)

**Cluster 8** D14 Post-vaccine 3 vs D0 Baseline

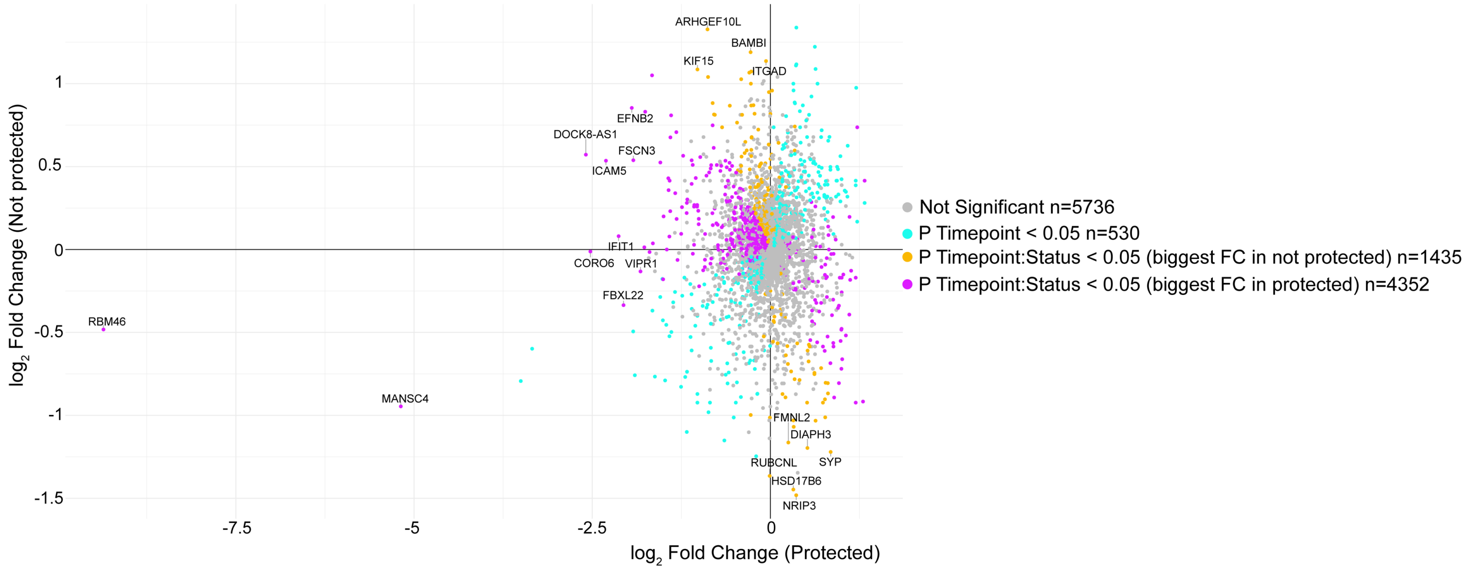


**Cluster 8** D5/6 Post-CHMI vs D0 Baseline

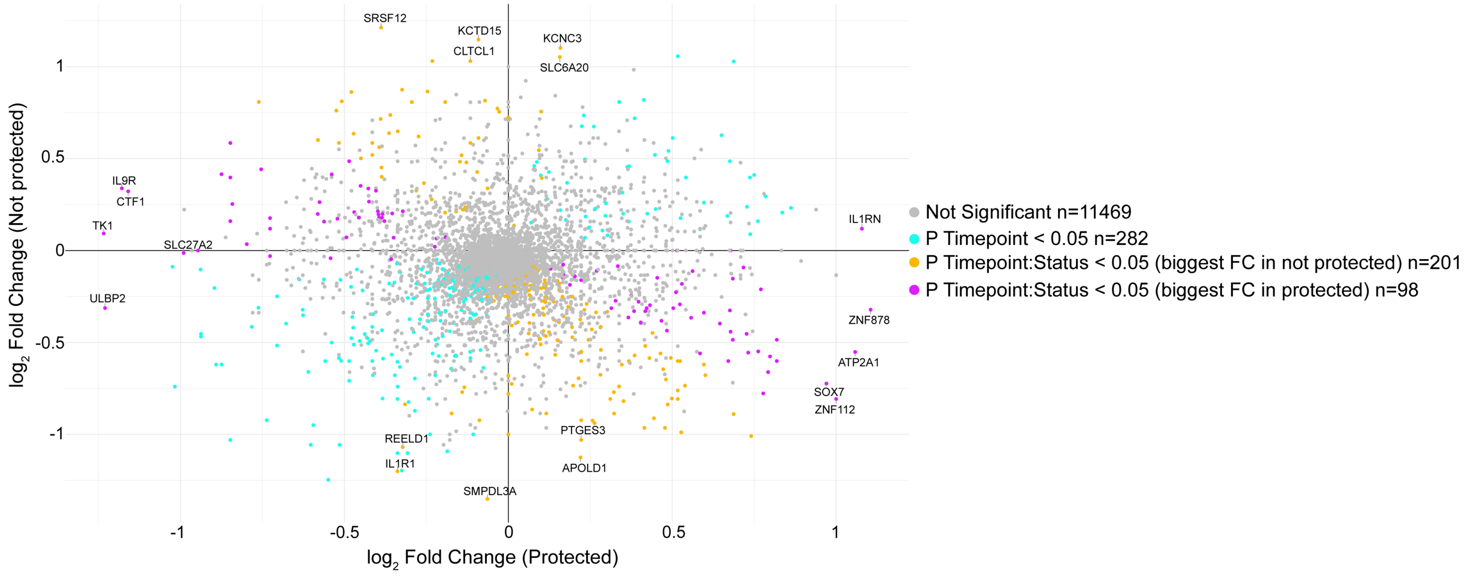


**Cluster 8** D5/6 Post-CHMI vs D14 Post-vaccine 3

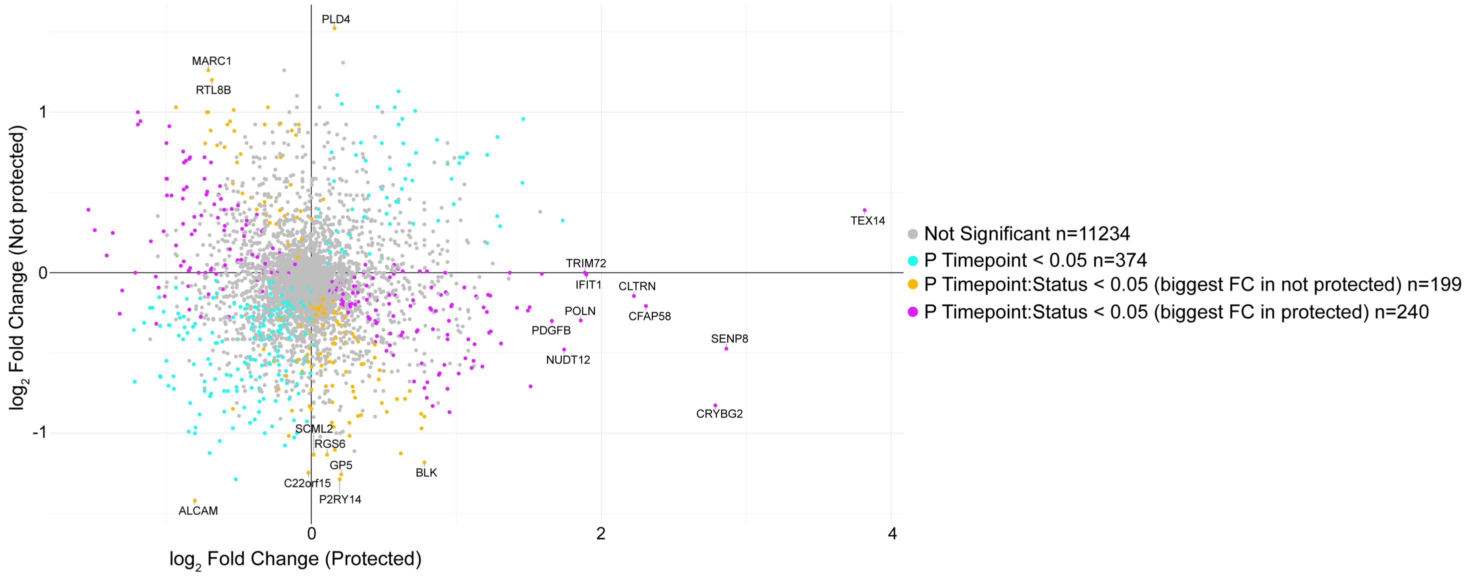

### Figure S8 (continued)

**Cluster 12** D14 Post-vaccine 3 vs D0 Baseline

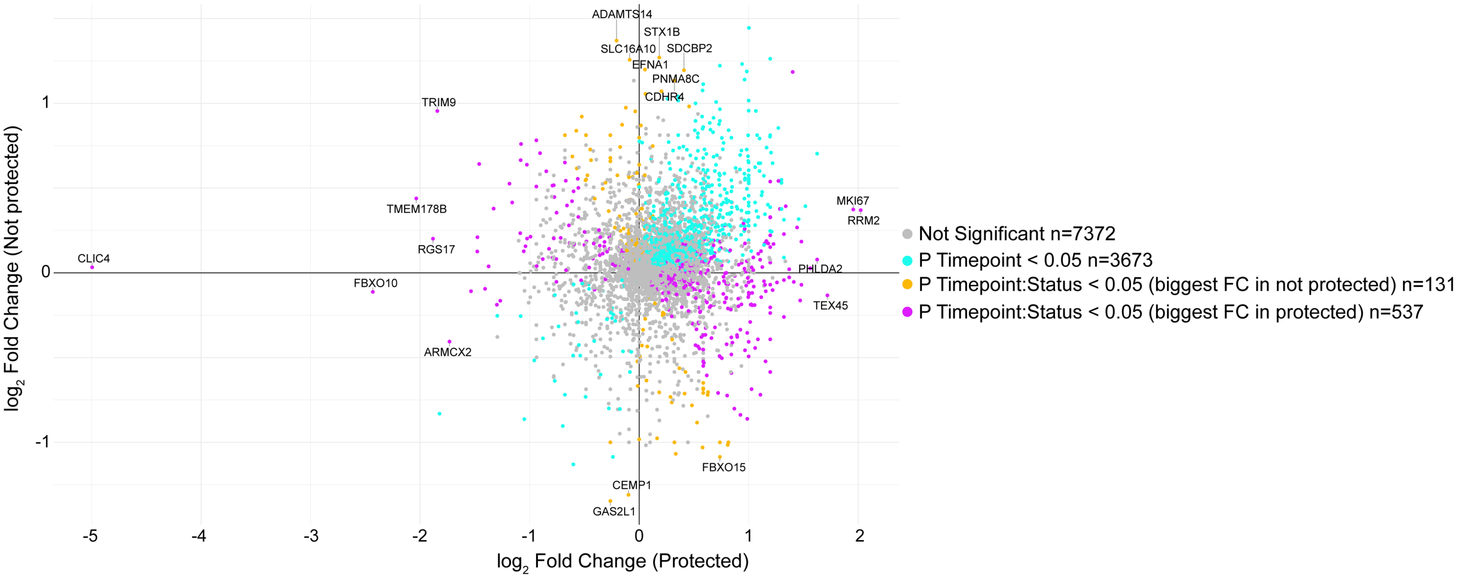


**Cluster 12** D5/6 Post-CHMI vs D0 Baseline

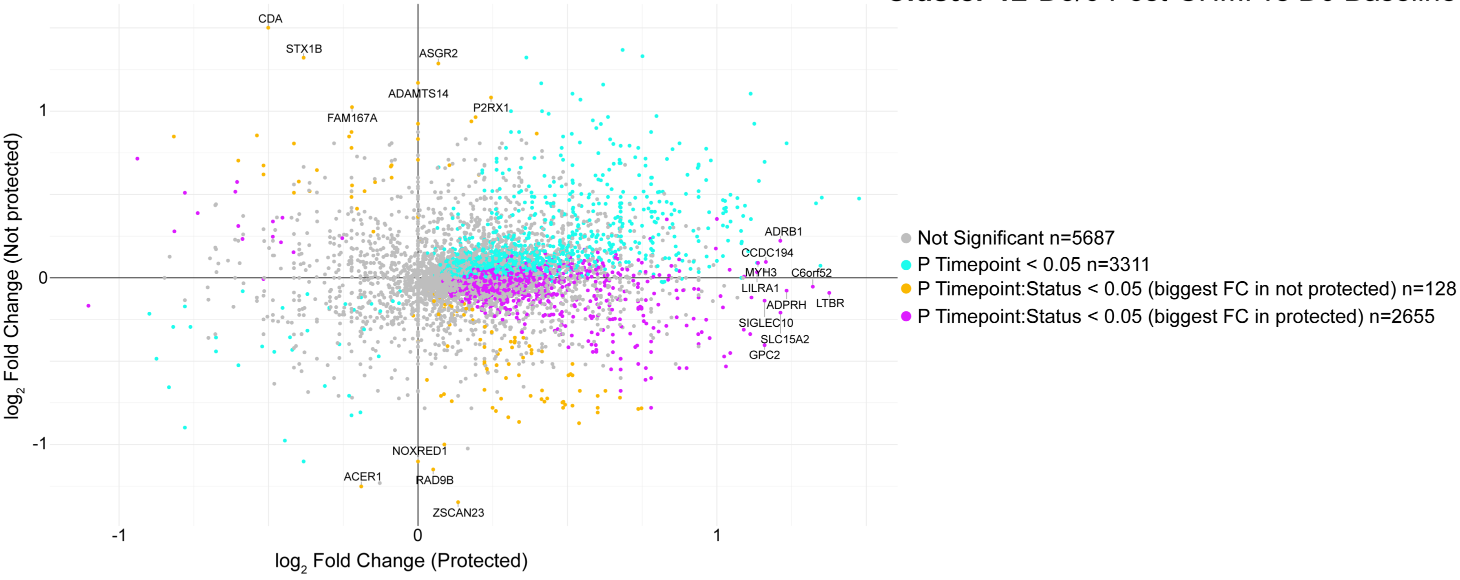


**Cluster 12** D5/6 Post-CHMI vs D14 Post-vaccine 3

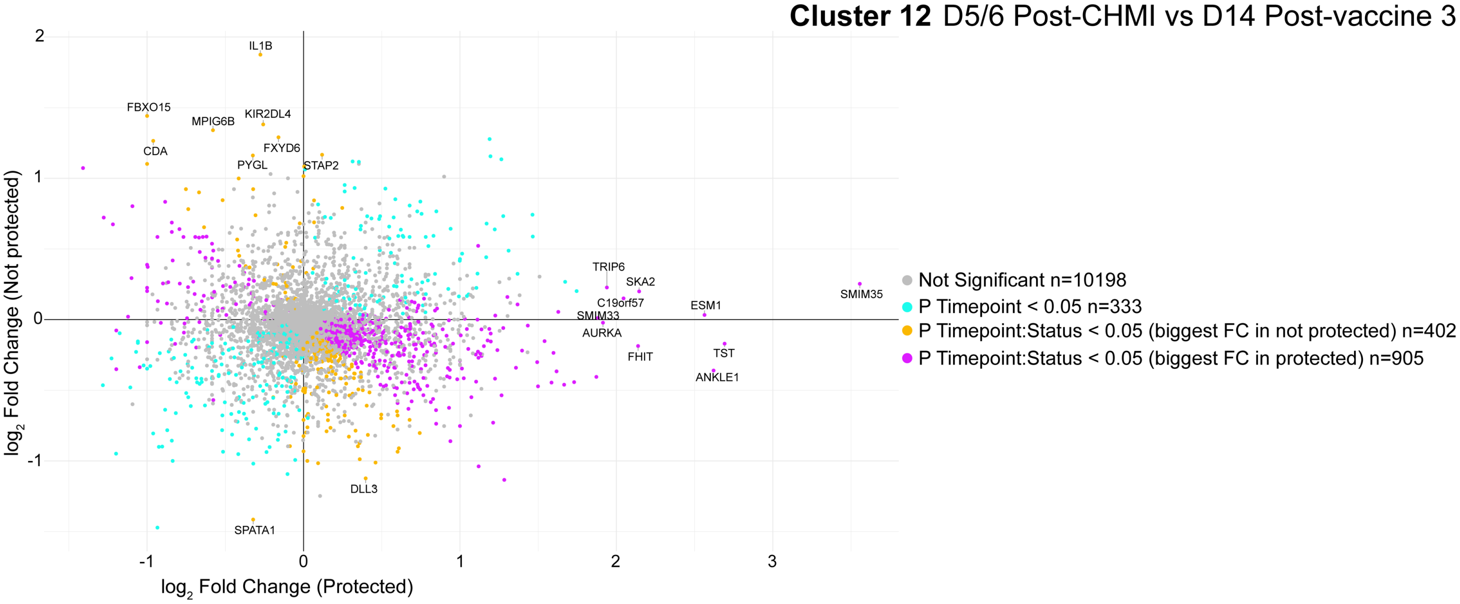

#### Figure S8 (continued)

**Cluster 18** D14 Post-vaccine 3 vs D0 Baseline

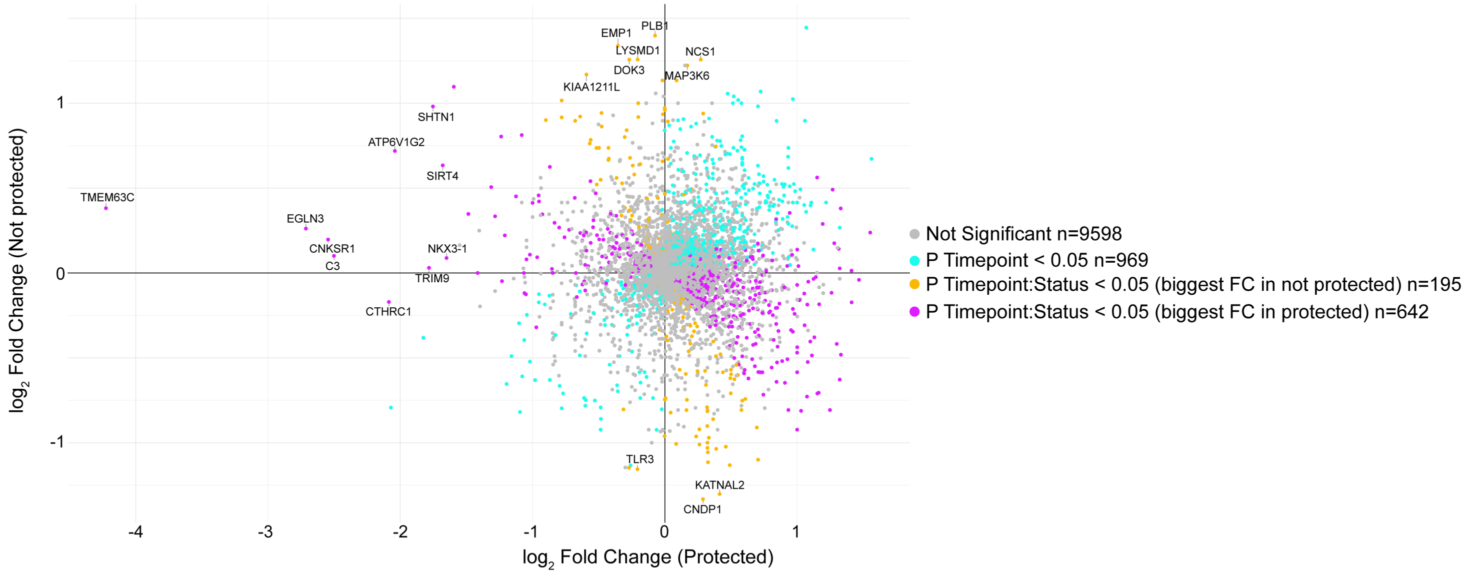


**Cluster 18** D5/6 Post-CHMI vs D0 Baseline

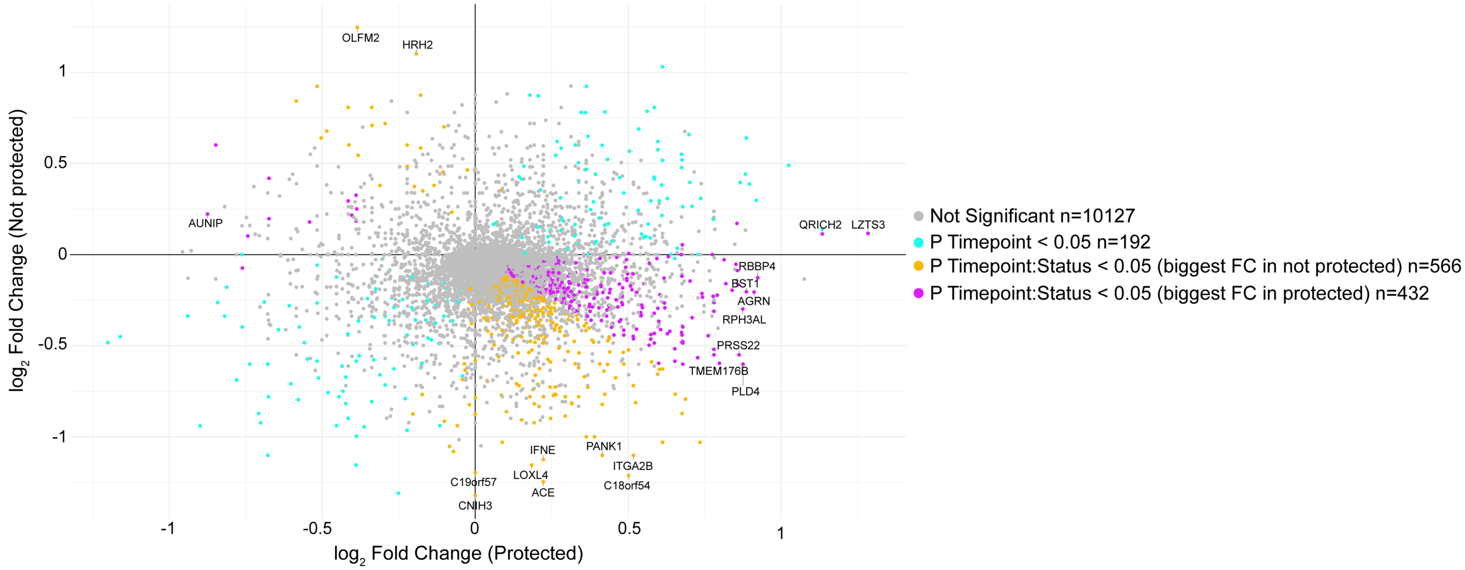


**Cluster 18** D5/6 Post-CHMI vs D14 Post-vaccine 3

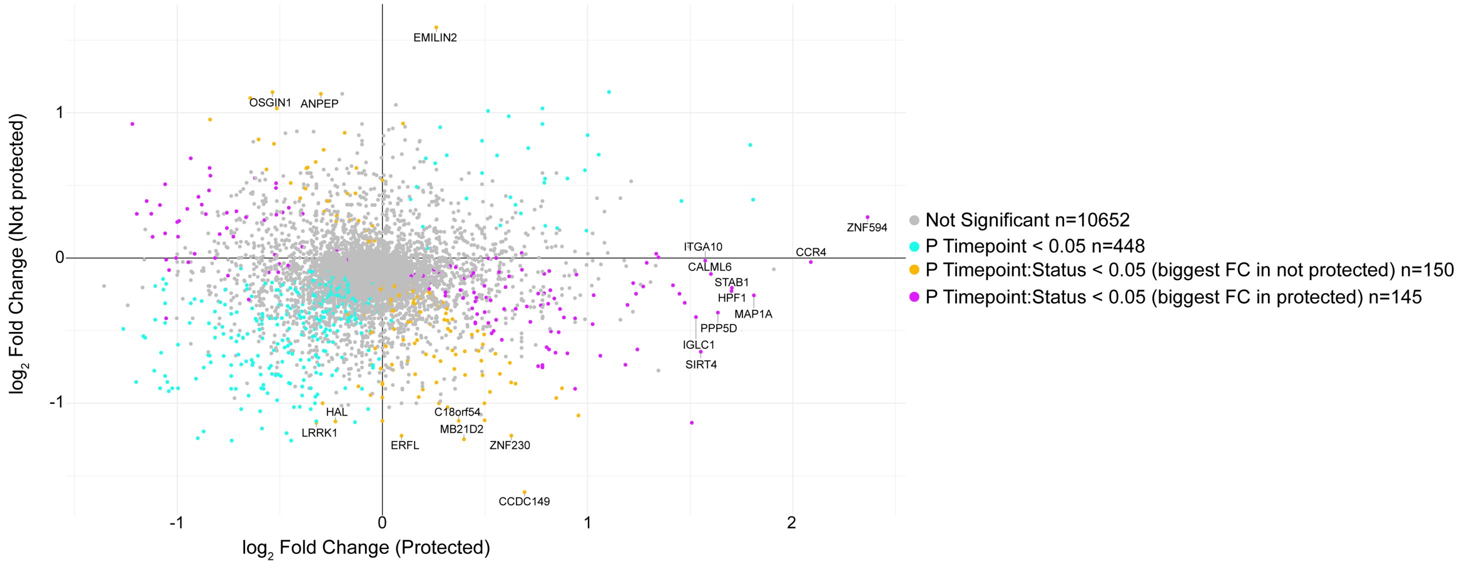

### Figure S8 (continued)

**Cluster 23** D14 Post-vaccine 3 vs D0 Baseline

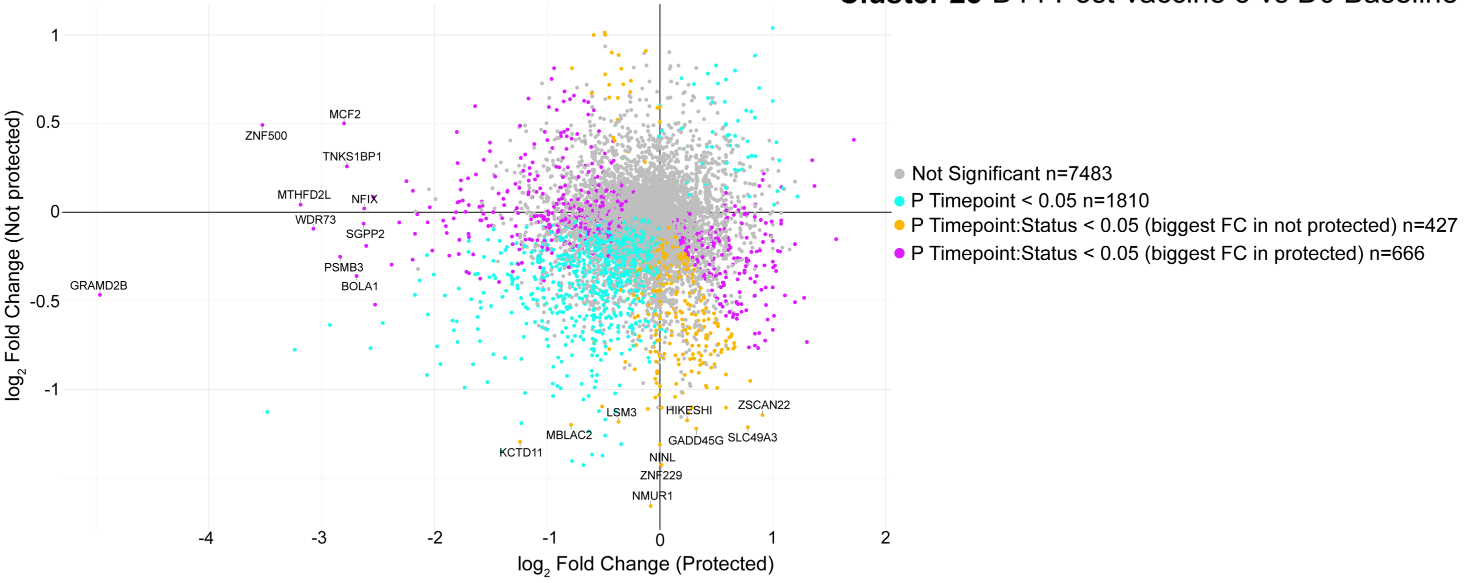


**Cluster 23** D5/6 Post-CHMI vs D0 Baseline

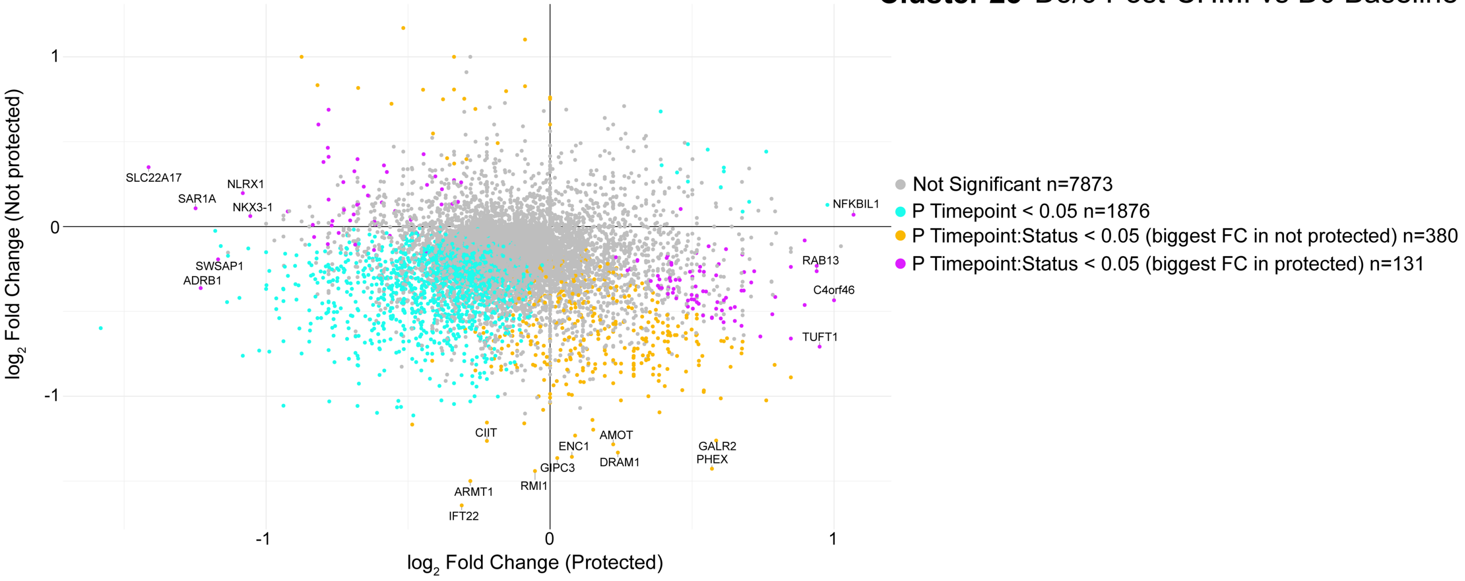


**Cluster 23** D5/6 Post-CHMI vs D14 Post-vaccine 3

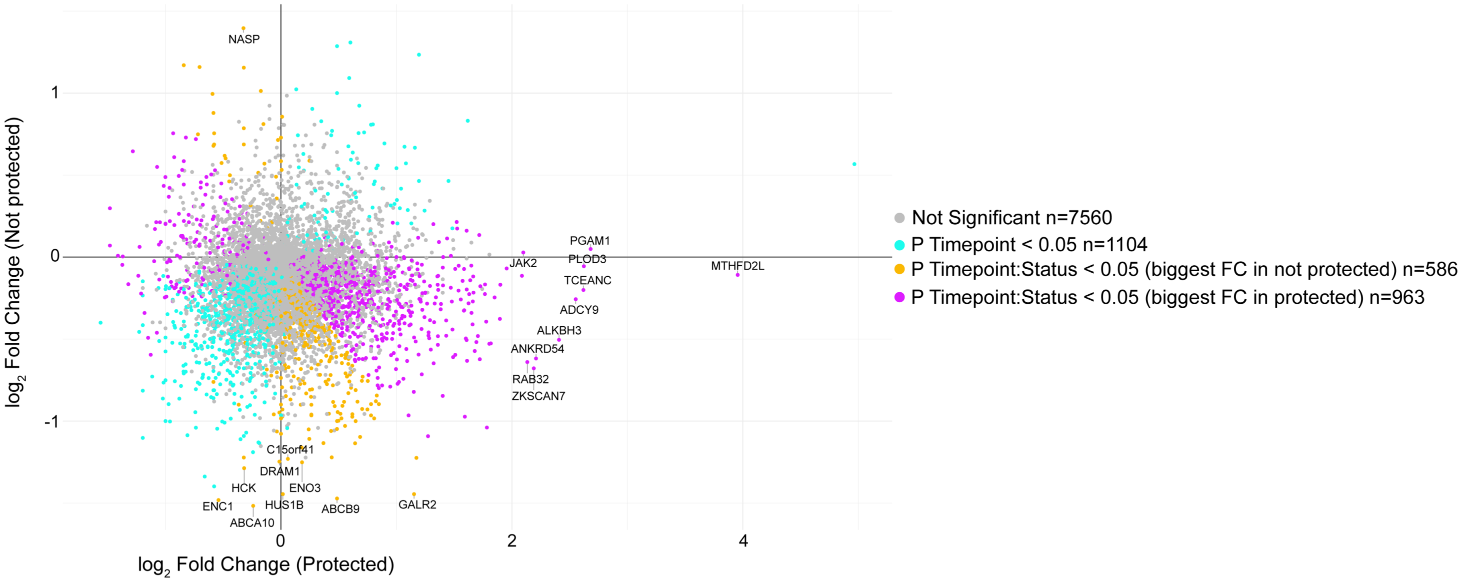

#### Figure S8 (continued)

**Cluster 24** D14 Post-vaccine 3 vs D0 Baseline

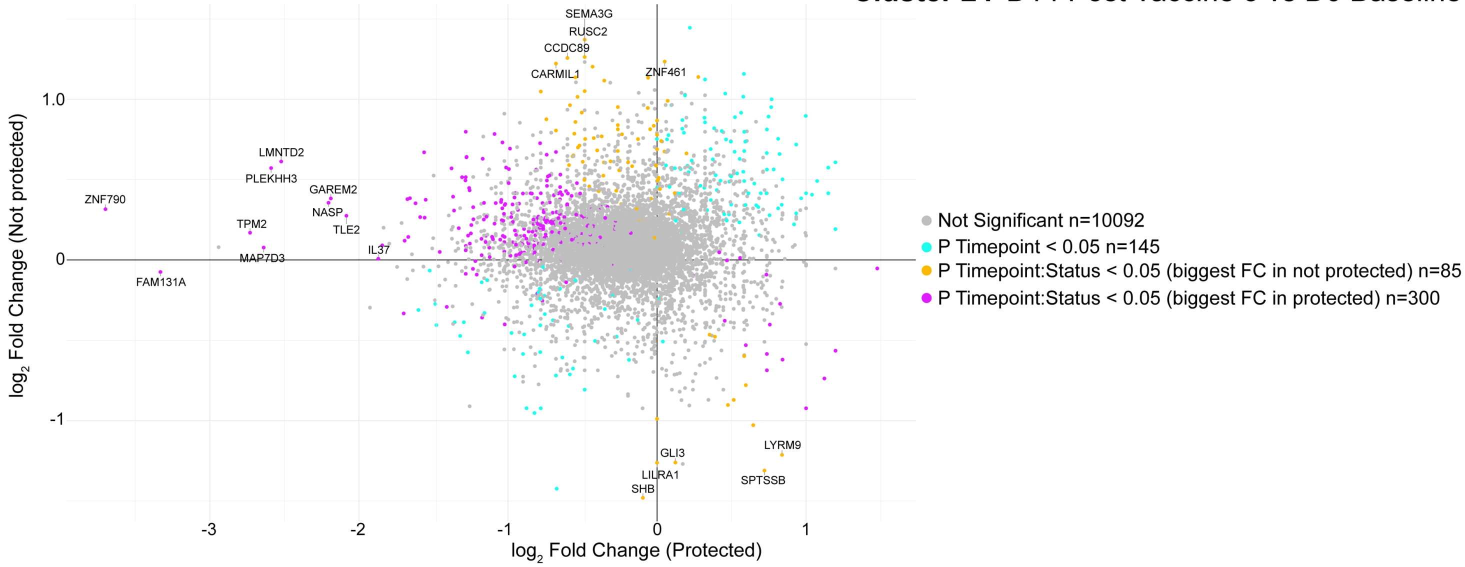


**Cluster 24** D5/6 Post-CHMI vs D0 Baseline

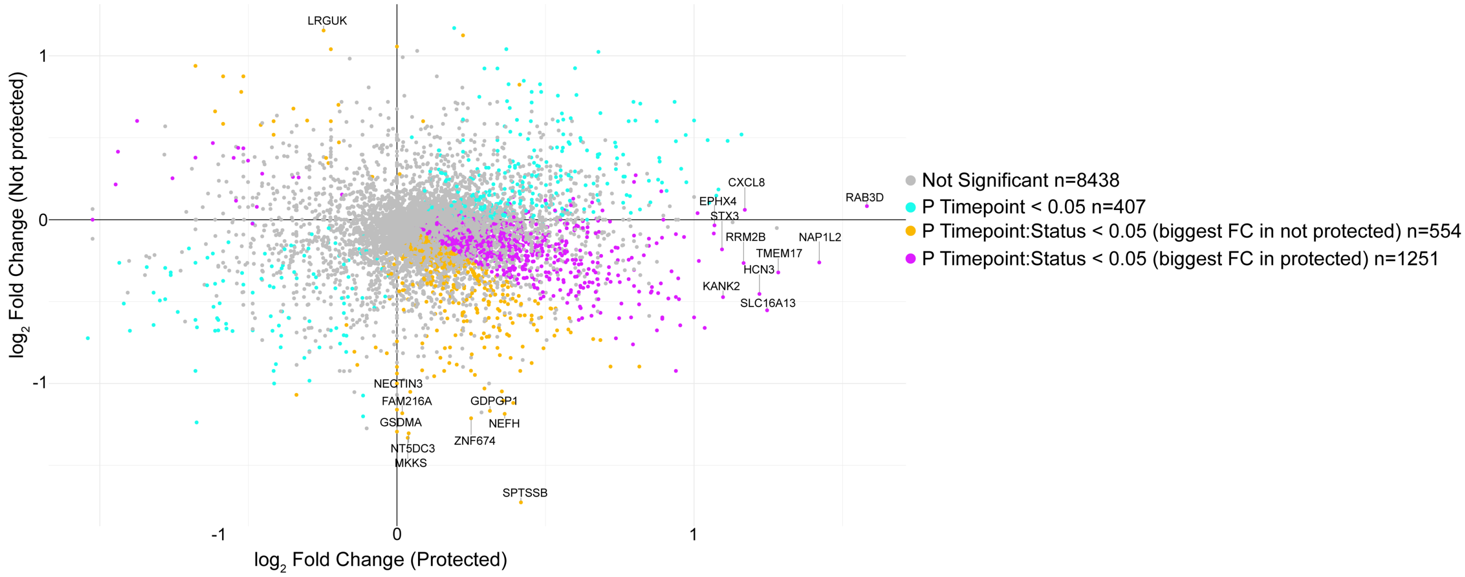


**Cluster 24** D5/6 Post-CHMI vs D14 Post-vaccine 3

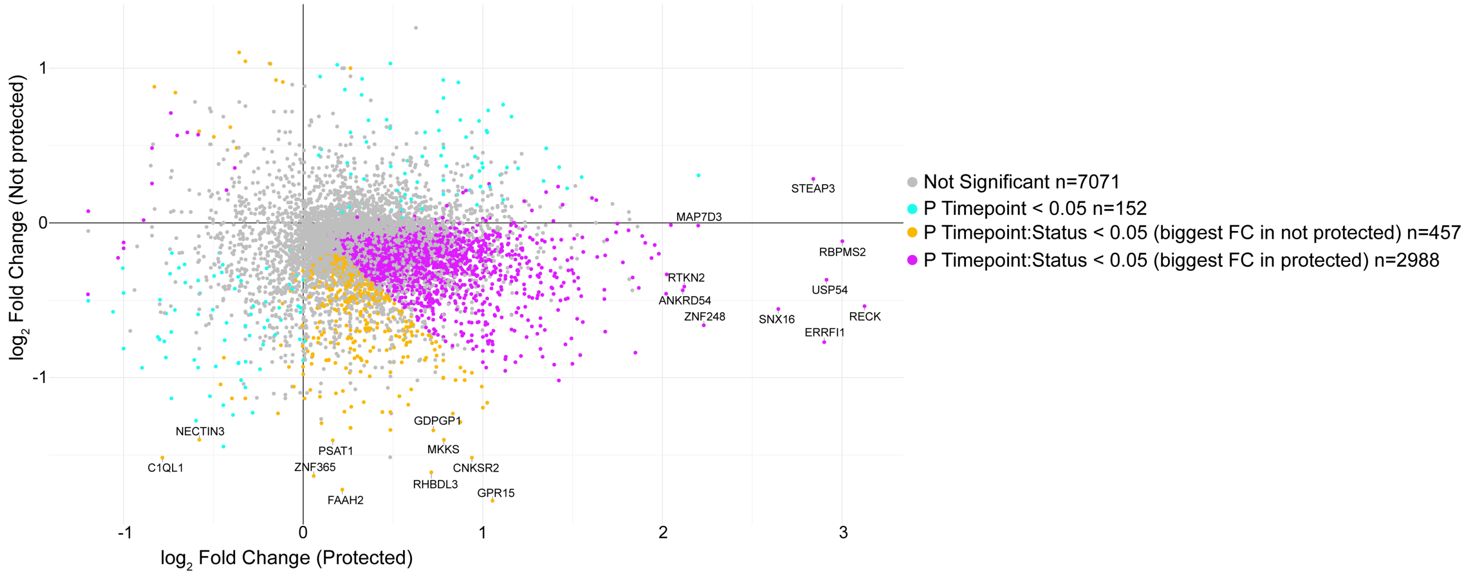

#### Figure S9: Frequency of TCRγδ+ cells, Vδ2+ cells, and Vδ2- cells in IMRAS participants

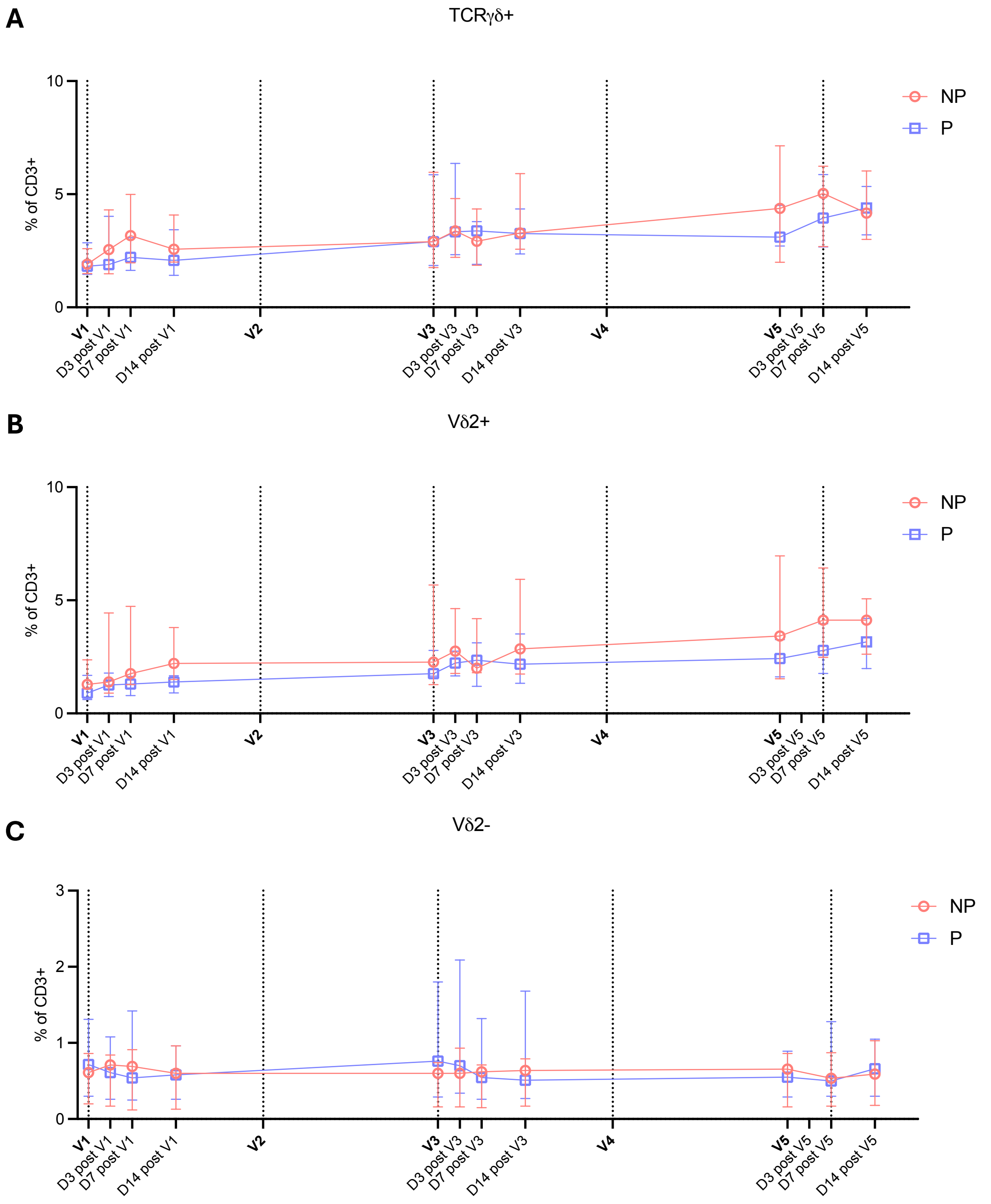

#### Figure S10: Total numbers of γδ T cells in the liver at day 4 post-immunization

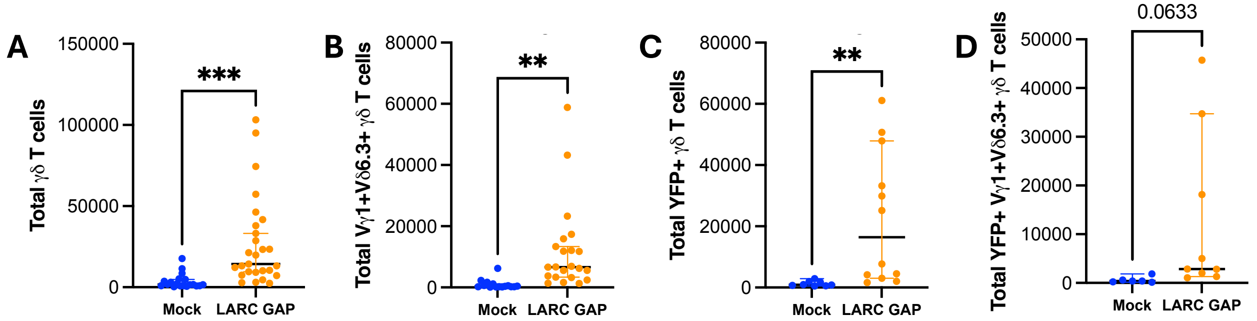

#### Figure S11: Flow cytometry gating strategies

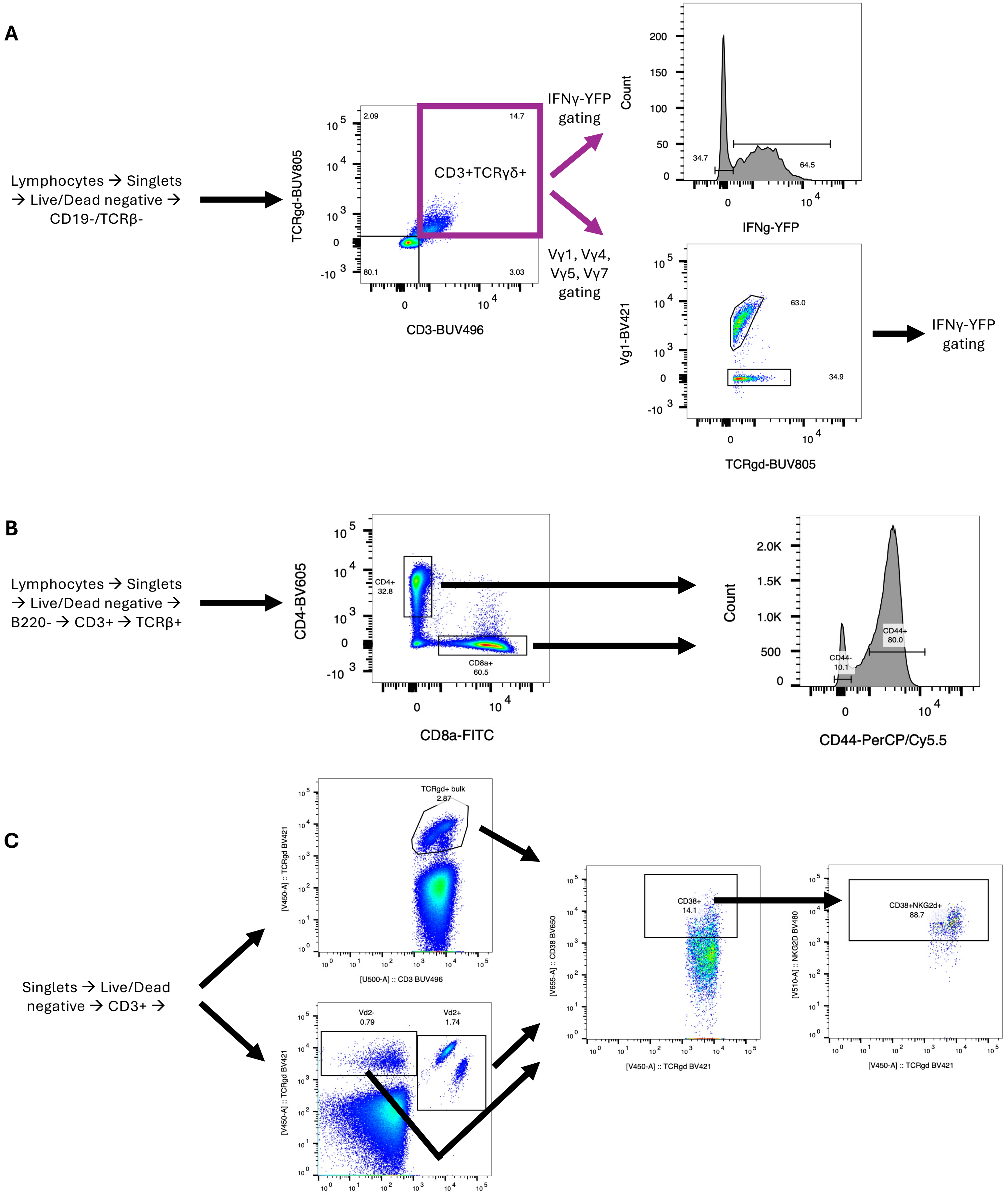
