## Supplemental Tables for "Hepatic γδ NKT cells modulate liver-resident CD8+ T cells to attenuated malaria parasite vaccines"

| Cluster | Protein markers |
|---|---|
| 1 | 2B4, PDL1, Sca1, Ly108, CD44 |
| 2 | 2B4, CD122, PDL1, CD38, Ly108, CD44 |
| 3 | Sca1, 2B4, Ly6C, PDL1, Ly108, CD44 |
| 4 | 2B4, CD44, CXCR6, Tigit, CD122, PDL1, CD38, CD39, Ly108 |
| 5 | PDL1, Sca1, Ly108, CD44 |
| 6 | PD1, PDL1, CD38, CD39, Sca1, Ly108, CD44 |
| 7 | Sca1, CD44, CXCR6, CD127, Tigit, PDL1 |
| 8 | Sca1, Ly6C, PDL1, Ly108, CD44 |
| 9 | Ly6C, PDL1, Ly108, CD44 |
| 10 | CD39, Sca1, CD44, 2B4, Ly6G, Ly6C, PDL1, CD38, Ly108 |
| 11 | Ly108, CXCR5, PDL1, CD38, Sca1, CD44 |
| 12 | Ly6C, 2B4, CD62L, PDL1, Sca1, Ly108, CD44 |
| 13 | Sca1, Ly108, CD44, PDL1 |
| 14 | Ly6C, Sca1, Ly108, CD44, 2B4, CD38, CD39 |
| 15 | CXCR6, Sca1, CD44, 2B4, Tigit, Ly6G, PDL1, CD38, CD39, Ly108 |
| 16 | Ly108, CD25, PDL1, CD38, CD39, Sca1, CD44 |
| 17 | Ly6C, Sca1, CD44, CXCR6, 2B4, Tigit, Ly6G, PDL1, CD38, CD39, Ly108 |
| 18 | Sca1, PD1, Ly6C, PDL1, Ly108, CD44 |
| 19 | Ly6C, CD39, Sca1, PD1, PDL1, CD38, Ly108, CD44 |
| 20 | Sca1, Tigit, CD62L, Ly6C, PDL1, CD39, Ly108, CD44 |
| 21 | KLRG1, Ly6C, Sca1, CD44, 2B4, PDL1, CD38, CD39, Ly108 |
| 22 | Ly6C, Sca1, CD44, CXCR6, Tigit, CD122, CD62L, Tim3, PDL1, CD38, CD39, Ly108 |
| 23 | CD44, 2B4, Ly6G, PDL1, CD38, CD39, Sca1, Ly108 |
| 24 | Sca1, Ly108, CD44, PD1, Tigit, Ly6G, CD122, CD62L, Tim3, CD69, PDL1, CD38, CD39 |
| 25 | CXCR6, 2B4, PD1, CD44, Lag3, PDL1, CD38, CD39, Ly108 |
| 26 | CXCR6, 2B4, CD122, PDL1, CD38, CD39, CD44, Tigit, Tim3, CD69, Sca1 |
| 27 | 2B4, CD122, CD44, Tigit, Ly6G, CD62L, Tim3, CD69, PDL1, CD38, CD39, Sca1, Ly108 |
| 28 | CD44, Tigit, Ly6G, CD!22, CD62L, Tim3, CD69, PDL1, CD38, CD39, Sca1 |

**Table S1: CyTOF γδ T cell clusters**

| Timepoint | Organ | Clusters* |
|---|---|---|
| 4 hours | Liver | 1 |
| Day 3 | Liver | 1, 20 |
| Day 7 | Liver | 1, 2, 3, 4, 5, 7, 10, 11, 12, 23 |
| Day 14 | Liver | 1, 13 |
| Day 3 | LDLN | 5, 8, 9 |
| Day 7 | LDLN | 9 |
| Day 3 | Spleen | 1, 3, 5, 6, 7, 8, 12, 13, 15, 16, 17 |
| Day 7 | Spleen | 1, 2, 5, 6, 8, 9 |
| Day 14 | Spleen | 1 |

* Table only shows clusters with average frequency of at least 1% that were enriched in mock-immunized mice compared to LARC GAP-immunized mice.

**Table S2: CyTOF clusters enriched in mock-immunized mice compared to LARC GAP-immunized mice**

| Target | Fluorophore | Clone | Source |
|---|---|---|---|
| Brilliant Stain Buffer | n/a | n/a | BD Biosciences |
| LIVE/DEAD Fixable Blue | n/a | n/a | Invitrogen |
| TCRβ | AF700 | H57-597 | Biolegend |
| CD19 | AF700 | 6D5 | Biolegend |
| CD3 | BUV496 | 17A2 | BD Biosciences |
| CD3 | BUV737 | 17A2 | BD Biosciences |
| TCRγδ | BUV805 | GL3 | BD Biosciences |
| Vγ1 | BV421 | 2.11 | Biolegend |
| Vγ4 | BV605 | UC3-10A6 | BD Biosciences |
| Vγ5 | APC | 536 | Biolegend |
| Vγ6 | PE | 1C10-1F7 | BD Biosciences |
| Vγ7 | BV711 | F2.67 | BD Biosciences |
| Vδ6.3 | BUV563 | 8F4H7B7 | BD Biosciences |
| CD4 | APC/Cy7 | GK1.5 | Biolegend |
| CD4 | BV605 | GK1.5 | Biolegend |
| CD8 | FITC | KT15 | MBL Life Sciences |
| CD8a | BUV737 | 53-6.7 | BD Biosciences |
| CD27 | BUV737 | LG.3A10 | BD Biosciences |
| CD44 | APC/Cy7 | IM7 | Biolegend |
| CD44 | PerCP/Cy5.5 | IM7 | Biolegend |
| CD62L | BUV563 | MEL-14 | BD Biosciences |
| CD69 | BUV395 | H1.2F3 | BD Biosciences |
| CD69 | BV421 | H1.2F3 | Biolegend |
| CD122 | PerCP/Cy5.5 | TM-B1 | Biolegend |
| CD127 | PE/Dazzle 594 | A7R34 | Biolegend |
| CD127 | APC | A7R34 | Biolegend |
| CD160 | PE/Cy7 | 7H1 | Biolegend |
| CCR6 | PE | 29-2L17 | Biolegend |
| ICOS | BV785 | C398.4A | Biolegend |
| KLRG1 | BV785 | 2F1/KLRG1 | Biolegend |
| PD1 | FITC | 29F.1A12 | Biolegend |
| B220 | APC/Fire750 | RA3-6B2 | Biolegend |
| IFNγ | PE/Cy7 | XMG1.2 | Biolegend |
| IL-17 | BV785 | TC11-18H10.1 | Biolegend |
| IL-4 | PE/Dazzle 594 | 11B11 | Biolegend |
| Ki67 | PerCP/Cy5.5 | 11F6 | Biolegend |

**Table S3: Flow cytometry antibodies**

| Target | Metal | Mass | Clone | Source |
|---|---|---|---|---|
| CD45 | y | 89 | 30-F11 | Standard Biotools |
| GR1 | Cd | 113 | RB6-8C5 | Biolegend |
| CD19 | Cd | 114 | 6D5 | Biolegend |
| MHCII | Cd | 116 | M5/114.15.2 | Biolegend |
| CD44 | Pr | 141 | IM7 | Biolegend |
| CD39 | Nd | 142 | Duha59 | Fluidigm |
| CD127 | Nd | 143 | A7R34 | Biolegend |
| CD38 | Nd | 144 | S21016C | Biolegend |
| CD25 | Nd | 145 | PC61 | Biolegend |
| CD8a | Nd | 146 | 53-6.7 | Biolegend |
| Ly6C | Sm | 147 | HK1.4 | Biolegend |
| TCRβ | Nd | 148 | H57-597 | Biolegend |
| CD4 | Sm | 149 | RM4-5 | Biolegend |
| TCRγδ | Sm | 150 | GL3 | Biolegend |
| CD62L | Eu | 151 | MEL-14 | Biolegend |
| KLRG1 | Sm | 152 | 2F1 | Biolegend |
| CD11b | Eu | 153 | ICRF44 | Biolegend |
| BST2 | Gd | 156 | 927 | Biolegend |
| CD11c | Gd | 157 | N418 | Biolegend |
| 4-1BB | Gd | 158 | 17B5 | Biolegend |
| CD122 | Tb | 159 | TM-β1 | Biolegend |
| CD69 | Dy | 160 | H1.2F3 | Biolegend |
| GITR | Dy | 161 | DTA-1 | eBioscience |
| PD-1 | Dy | 162 | 29F.1A12 | Biolegend |
| CD73 | Dy | 163 | TY/11.8 | Biolegend |
| CXCR5 | Dy | 164 | L138D7 | Biolegend |
| Ly6G | Er | 166 | 1A8 | Biolegend |
| CXCR6 (CD186) | Er | 167 | SA0151D1 | Biolegend |
| PDL1 | Er | 168 | 10F.9G2 | Biolegend |
| Tim3 | Tm | 169 | RMT3-23 | Biolegend |
| Ly108 | Yb | 170 | 13G3 | Biolegend |
| Tigit | Yb | 171 | 1G9 | Biolegend |
| Sca-1 | Yb | 172 | E13-161.7 | Biolegend |
| 2B4 | Yb | 173 | m2B4 (B6)458.1 | Biolegend |
| Lag3 | Yb | 174 | C9B7W | Biolegend |
| Siglec-F | Lu | 175 | S17007L | Biolegend |
| NK1.1 | Yb | 176 | S17016D | Biolegend |

**Table S4: Mass cytometry antibodies**

| Vδ1 module features | Vδ2 module features | Cytotoxic/Cytolytic features | IFNG-signaling features |
|---|---|---|---|
| KLRF1 | KLRC1 | KLRK1 | IFNG |
| FCGR3A | IL7R | KLRF1 | IL2RB |
| CCL3 | NOSIP | KLRC1 | TBX21 |
| TRGC2 | TRGC1 | GZMB | GATA3 |
| CMC1 | LTB | GZMH | ITGB1 |
| GZMB | GZMK | GZMK | STAT1 |
| FGFBP2 | ALOX5AP | GNLY | IRF1 |
| TYROBP | TRAC | NKG7 | IRF4 |
| GZMH | JUN | PRF1 | IRF8 |
| CD63 | CD44 | CCL4 | KLRB1 |
| PLAC8 | NFKBIA | CCL5 | IFNGR1 |
| GNLY | CD52 | | EOMES |
| PRF1 | ZFP36L2 | | ICOS |
| TRBC2 | | | |
| GSTP1 | | | |
| NKG7 | | | |
| C12orf75 | | | |
| HOPX | | | |
| CCL4 | | | |
| SUB1 | | | |
| IFITM1 | | | |
| KLF2 | | | |
| EFHD2 | | | |
| IFITM2 | | | |

**Table S5: Genes used to define Vδ1, Vδ2, cytotoxic/cytolytic, and IFNγ-signaling modules.**
